# Effects of protein interface mutations on protein quality and affinity

**DOI:** 10.64898/2026.03.24.713863

**Authors:** Jurrian K. de Kanter, Eva Smorodina, Aygul Minnegalieva, Marloes Arts, Lasse Møller Blaabjerg, Marina Frolenkova, Puneet Rawat, Lina Wolfram, Helena Britze, Yano Wilke, Lucas Weissenborn, Laurens Lindenburg, Emily Engelhart, Kerry L. McGowan, Ryan Emerson, Randolph Lopez, Joke Gerarda van Bemmel, Samuel Demharter, Roberto Spreafico, Victor Greiff

## Abstract

Accurately modeling antibody-antigen interactions requires distinguishing intrinsic binding affinity (“*protein-interaction*”) from protein biophysical properties (“*protein-quality*”), including folding, stability, and expression. However, high-throughput mutational measurements commonly used to train and benchmark computational models often conflate these effects, obscuring the true determinants of molecular recognition. Here, we present an experimental and analytical framework to disentangle *protein-interaction* effects from *protein-quality* effects in single-domain antibody (VHH)-antigen binding. Using a large-scale deep mutational scanning (DMS) dataset spanning four VHH-antigen complexes, with single and double mutations in both partners, we introduce control binders to quantify *protein-quality* changes independently of *protein-interaction*. This enables decomposition of experimentally measured affinity into *protein-interaction* and *protein-quality* components at scale. Leveraging the disentangled dataset, we evaluated state-of-the-art structure- and sequence-based models for *protein-quality* and *protein-interaction* prediction and show that their performance largely reflects *protein-quality* rather than *protein-interaction* effects. Our results highlight a major confounder in current datasets and suggest that accounting for *protein-quality* will be essential for training next-generation affinity-prediction models.

**Nomenclature:** *Antibody related terms:*

- **Primary VHH**: The VHH of a VHH-antigen complex for which the paratope and the epitope weremutated.
- **Control VHH**: A second VHH that binds to the same antigen as the primary VHH but has non-overlapping epitope positions and therefore does not bind to any of the mutated antigen positions.

*Affinity-related terms:*

- **Real Affinity**: “The strength of the interaction between two […] molecules that bind reversibly (interact)” ^1^. In the context of antibody-antigen binding, it quantifies interactions between active proteins (which are expressed and correctly folded ^2^ and are therefore functionally and biologically active (see below). It is commonly quantified by the equilibrium dissociation constant, K_D_.
- **Observed affinity (°K_D_):** The interaction strength experimentally measured between two molecules. Unlike real affinity, this value is confounded by the biophysical properties of the individual binding partners, specifically their folding, stability, and expression levels. Consequently, the observed affinity often differs from the real/intrinsic affinity if a significant fraction of the protein population is inactive ^3^. **NOTE: Unless otherwise specified, °K_D_ is reported in - log10 space. For example, a °K_D_ of -9 corresponds to 10^-9^M or 1nM.**
- **Change in observed affinity (Δ°K_D_)**: The shift in the observed affinity between two proteins upon mutation, reported as the log_10_-transformed fold change. A value of 1 reflects a 10-fold difference, a value of 2 a 100-fold difference, etc. This aggregate change resolves into two distinct biophysical components ^2, 4^:

- ***Protein-interaction* change:** The change in the intrinsic thermodynamic affinity between the two binding partners, each in its active state (i.e., the specific change in interface Gibbs free energy because both enthalpy and entropy are considered).
- ***Protein-quality* change:** The change in the fraction of the mutated protein population that is biologically active - meaning it is expressed, correctly folded, and stable ^2, 5^.

- **Folding:** The process that guides the polypeptide chain toward its native conformation, which is a prerequisite for forming a functional binding site.
- **Stability:** The thermodynamic capacity to maintain the folded structure over time and under physiological conditions. Stability (decrease in Gibbs free energy from the unfolded to the folded state) ensures the binding interface remains intact and prevents competing processes such as aggregation ^6^.
- **Expression:** The steady-state abundance of the protein. This is largely dependent on proper folding and stability, as cellular quality control mechanisms degrade proteins that fail to fold or remain stable at functional concentrations.
- **Change in relative affinity (ΔΔ°K_D_)**: the difference between the Δ°K_D_ of the primary VHH compared to the control VHH for a given epitope mutation.

*Model-related terms:*

- **ESM-IF1 sc**: Single-chain (sc) structure-conditioned inverse folding model (ESM-IF1), using the isolated monomer structure of the mutated protein: either the VHH or the antigen ^7^.
- **ESM-IF1 mc**: Multi-chain (mc) structure-conditioned model (ESM-IF1), using the full complex structure (both antibody and antigen) ^7^.
- **Stability prediction score**: Score that represents the predicted change in stability based on a single mutation, normally represented as ΔΔG.

## Introduction

The ability to predict the effects of mutations on protein-protein interactions is fundamental to understanding molecular recognition and engineering high-affinity binders ^8–10^. In antibody development, protein-protein interaction predictors are increasingly applied to assist rational design, accelerate therapeutic optimization, interpret immune repertoires and *de novo* design ^11–19^. Recently, deep learning-based structure and sequence models, including inverse folding, have shown great promise in in silico protein engineering and are reported to capture binding determinants ^20^. However, a critical question remains: to what extent do these methods learn patterns of *protein-interaction*, versus detecting *protein-quality* changes of the mutated protein ^21, 22^. Here, we define *protein-interaction* change as the change in the intrinsic thermodynamic affinity between the functional states of the two binding partners (i.e., the specific change in interface energy), and we define *protein-quality* change as the change in the fraction of the mutated protein population that is biologically active - meaning it is expressed, correctly folded, and stable.

Training and benchmarking protein-protein interaction models require massive datasets^23^. Recent studies have leveraged high-throughput assays such as deep mutational scanning to quantify the change in observed affinity of thousands of variants ^24–27^. Yet, interpreting these data is complicated by a biophysical confounder: mutations often induce *protein-quality* changes ^2, 28^, which affect the observed affinity, even when the real/intrinsic affinity remains unchanged. This conflation of a protein’s structural state with its interaction strength makes it impossible to correctly evaluate model performance on real affinity prediction on the mixed signal of observed affinity and hampers model training on real affinity.

Here, we address this challenge by systematically disentangling the effects of 7,185 mutations on *protein-qualit*y versus *protein-interaction* in antibody-antigen complexes. We investigated four complexes of a heavy chain-only antibody variable domain (VHH) and its antigen, introducing single and double mutations in both partners separately. Crucially, for each VHH-Ag complex, we utilized a control VHH binding to a non-overlapping epitope on the same antigen to control for antigen *protein quality,* thereby separating intrinsic affinity shifts from protein quality effects.

We used this dataset, where *protein-quality* and *protein-interaction* effects of mutations are separated, to benchmark how well existing inverse folding models can predict these two components. Our results reveal that mutant scores of current state-of-the-art models, such as log-likelihoods, are primarily associated with *protein-quality* changes, with only a minor association with interface-specific *protein-interaction* changes. Together with the continued expansion of curated antibody-antigen structural resources for computational modelling ^23, 29, 30^, these findings underscore the need for datasets and modelling approaches that explicitly separate *protein-interaction* from *protein-quality* effects, an essential step toward truly predictive models of antibody-antigen recognition.^29^

## Results

### Decomposing observed affinity of thousands of VHH-antigen mutant pairs into protein quality and protein interaction components

To comprehensively investigate antibody-antigen interactions, we separately assessed the affinity of epitope and paratope mutants using the yeast-display AlphaSeq assay (Fig. 1A, B) ^31^. In this assay, yeast mating efficiency is driven by the interaction of surface-displayed partners. Consequently, the observed affinity depends on both the intrinsic affinity of the complex and the surface abundance of the proteins^32^. Here, we differentiate between the real affinity (K_D_): “the strength of the interaction between two […] molecules that bind reversibly (interact)”^1^, and the observed affinity (°K_D_): the interaction strength experimentally measured, here by AlphaSeq, between two molecules. Unlike the real affinity, this observed affinity value is confounded by the biophysical properties (*protein-quality,* see Nomenclature) of the individual binding partners (Fig. 1A). Consequently, the observed affinity often differs from the real/intrinsic affinity if a significant fraction of the protein population is inactive. We selected four nanomolar-affinity VHH-antigen pairs for which experimental structures were available. This allowed us to select control VHHs with non-overlapping epitopes, and to assess biophysical properties of mutations (see below). The selected VHHs had antigens SARS-CoV-2 RBD (PDB: 7olz, 7z1b) and Botulinum neurotoxin (PDB: 7m1h, two distinct VHHs) (Table 1).

**Figure 1.**
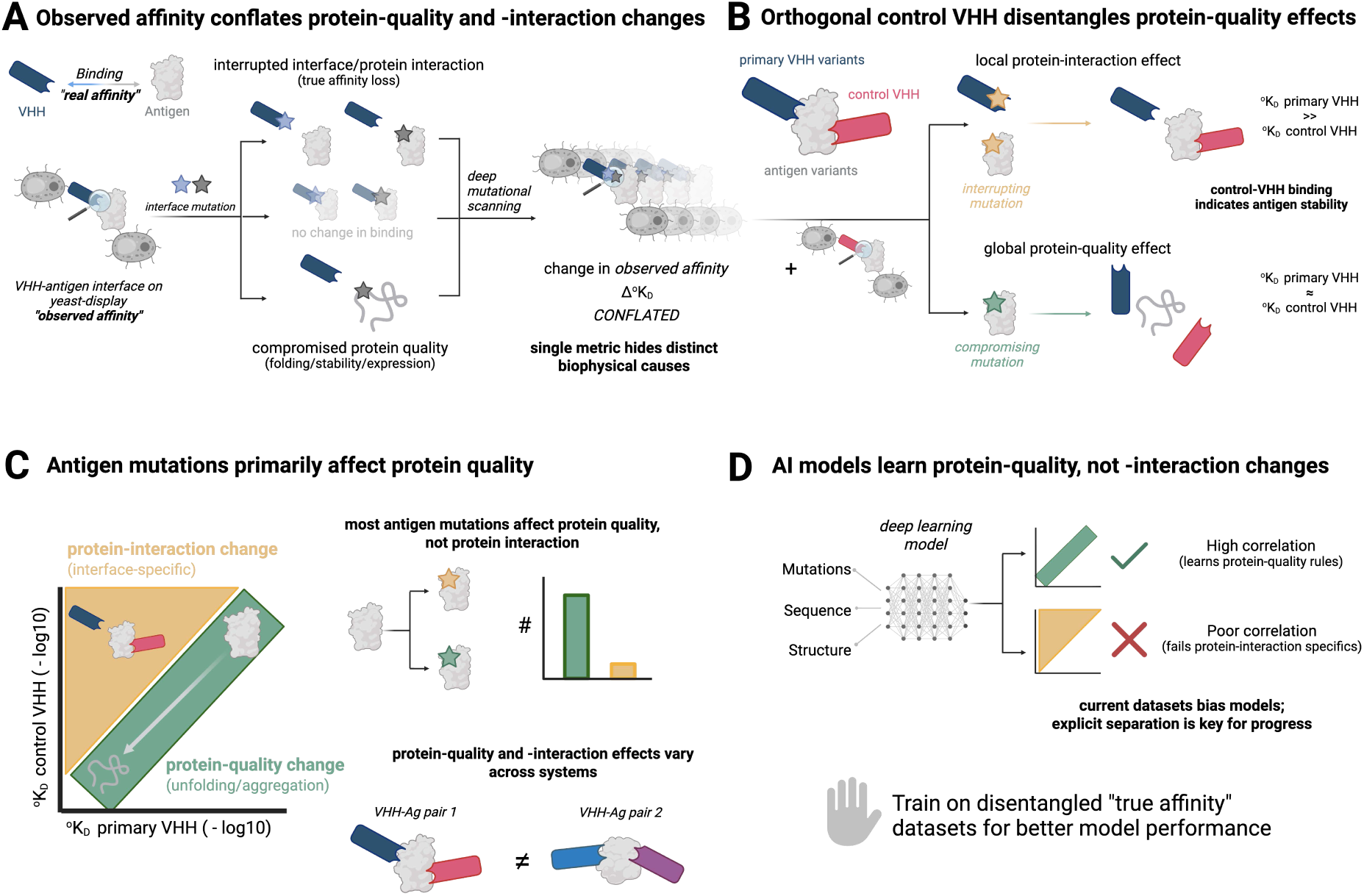
Protein-quality and protein-interaction components of observed affinity of deep mutational scanning of VHH-antigen complexes and using it to benchmark inverse folding (-derived) models. A) Mutations can change antibody-antigen recognition both by directly disrupting interface contacts (paratope-epitope interaction) and by altering overall protein quality (folding, stability, expression). Separating these effects is critical for understanding and calculating real affinity. B) We present deep mutational scanning data for VHH and antigen variants. We monitor antigen protein quality through orthogonal binding of a control VHH. C) Correlating observed primary and control K_D_ values enables us to distinguish between mutations that affect protein interaction vs protein quality. We show that across VHH-antigen complexes, most antigen substitutions reduce binding primarily by degrading antigen protein quality rather than disrupting interface energetics; most antigen positions contribute to protein-quality effects while few are true protein-interaction positions. The extent of these effects varies between complexes. D) Existing deep learning models (structure-conditioned (inverse-folding) models, free-energy predictors) predominantly capture protein-quality effects of antigen mutations. We conclude that reporting real affinities adjusted for protein quality changes in future (training) datasets is desirable.

**Table 1.** VHH target-antigen-VHH control complexes used in this study.

| Complex name | UniProt (ID) | Antigen (name, PDB, chain) | Primary VHH (PDB, chain) | Main control VHH (PDB, chain) | Additional control VHHS (PDB, chain) | WT resolution (primary and main control VHH-antigen) (Å) | Experimental method | AlphaSeq primary VHH WT $^oK_D$ (nM) | AlphaSeq main control VHH $^oK_D$ (nM) |
| --- | --- | --- | --- | --- | --- | --- | --- | --- | --- |
| P0DTC2-7olz_B | P0DTC2 | SARS-CoV-2 receptor-binding domain (RBD), A | 7olz, B | 8sk5, B | 7z1b, E<br>7x2l, B<br>7fh0, B | 1.75, 2.01 | X-ray, X-ray | 54nM | 201nM |
| P0DPI0-7m1h_C | P0DPI1 | Botulinum neurotoxin A (botA), A | 7m1h, C | 7m1h, G | 7m1h, B<br>7m1h, F | 2.78, 2.78 | X-ray, X-ray | 163nM | 100nM |
| P0DPI0-7m1h_G | P0DPI1 | Botulinum neurotoxin A (botA), A | 7m1h, G | 7m1h, C | 7m1h, F | 2.78, 2.78 | X-ray, X-ray | 69nM | 136nM |
| P0DTC2-7z1b_E | P0DTC2 | SARS-CoV-2 receptor-binding domain (RBD), B | 7z1b, E | 6waq, C | - | 2.30, 2.20 | X-ray, X-ray | 260nM | 87nM |

The paratope and epitope of each “primary VHH” were mutated (see Methods). To isolate *protein-quality* changes from *protein-interaction* changes, we included a “control VHH” for each complex that binds a non-overlapping epitope on the same antigen (Fig. 1B). Given that the control VHH binds a distinct antigenic site, any change in its observed affinity upon mutating the epitope of the primary VHH serves as a proxy for the change in *protein-quality* of the antigen variant. We introduced all possible single amino acid substitutions in the paratope and epitope of the primary VHH. In addition, a selection of double mutants within the paratope and epitope were included (see Methods). In total, we measured the change in observed affinity for 1,904 single and 1,858 double mutants in the antigens, and 1,257 single and 1,615 double mutants in the primary VHHs. The AlphaSeq assay measures all antigen variants vs all primary-VHH variants within a given complex. Here, we focused on the interactions for which *protein-quality* and *protein-interaction* could be separated in the experimental data, and those that allowed for the evaluation of existing (inverse folding) models that score the effect of single protein mutations on these two characteristics. These were the interactions of 1) the wild-type (WT) VHH with all variants of the antigen and 2) the WT antigen with all VHH variants, but *not* of the interaction between the VHH variants with the antigen variants.

### Antigen mutational scan reveals high contributions of *protein-quality* changes to observed affinity

First, we quantified the contribution of *protein-quality* and *protein-interaction* to the change in observed affinity upon mutating the antigen. Consistent with previous reports ^3, 26, 33^, 83.6%-93.2% of single mutations and 89.4%-98.9% of double mutations exhibited a lower observed affinity than the wild type across the four studied primary VHH-antigen pairs (Supp. Fig. 1A). When grouped by position, mutational tolerance varied significantly along the antigen, primary VHH, and control VHH (Fig. 2A, Supp. Fig. 1B-D).

**Figure 2.**
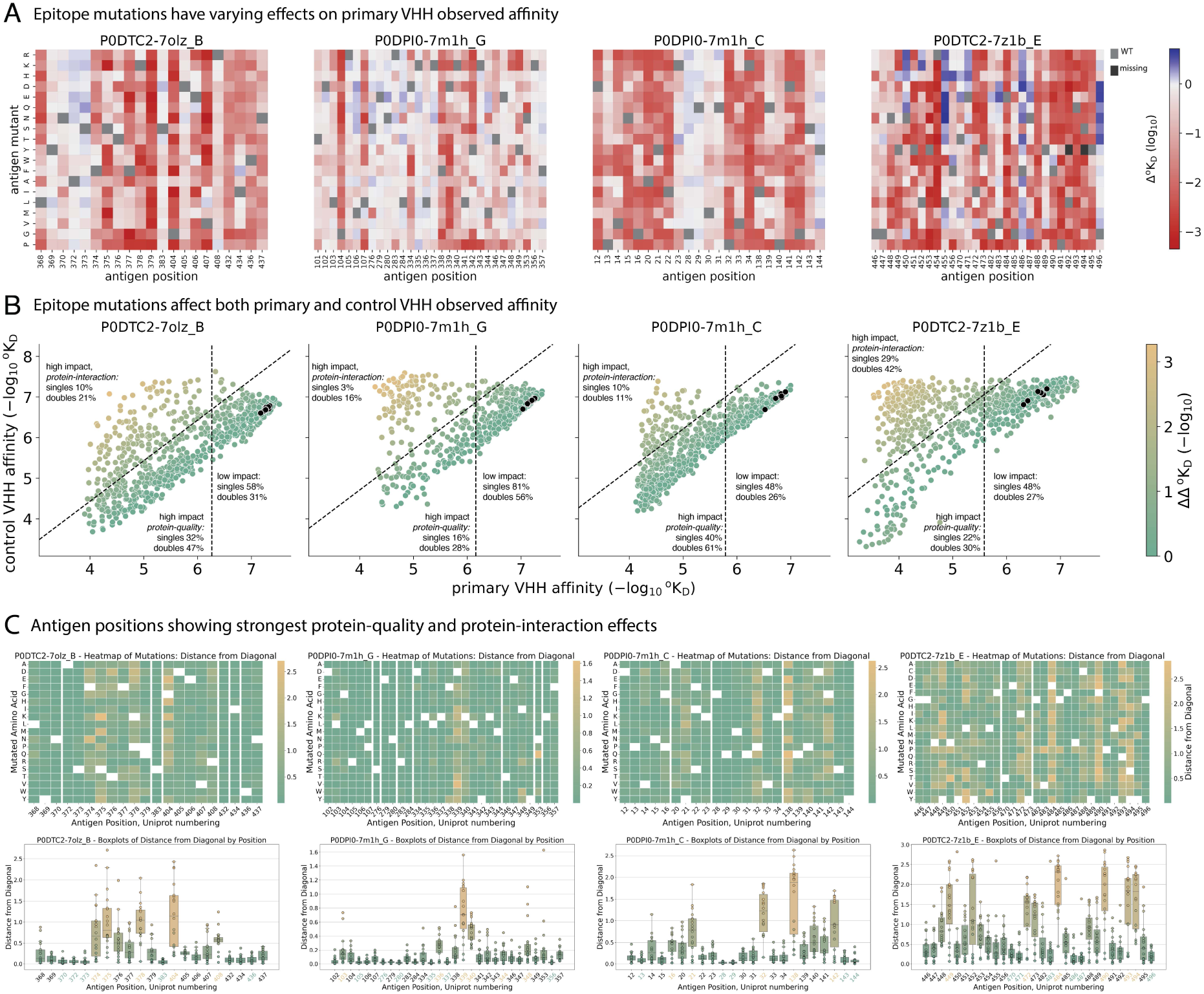
The majority of antigen mutations have a negative Δ°K_D,_ driven by decreased protein quality. (**A**) Antigen DMS heatmaps of structural epitope positions, primary VHH change in observed affinity (Δ°K_D_). Black signifies a missing measurement; grey signifies wild type. See Supp. Fig. 1B for DMS heatmaps of epitope positions for the control VHH (B) Primary VHH °K_D_ compared to control VHH °K_D_ of single epitope mutations, colored by ΔΔ°K_D_, i.e., the difference between the change in °K_D_ in the primary VHH compared to the control VHH. The higher the number on the x-axis, the stronger the affinity of the given antigen-mutant/parental-VHH pair. The higher the number on the y-axis, the higher the affinity of the given antigen-mutant/control VHH pair. In each panel, the following percentages of the mutations are indicated: “low impact” = Δ°K_D_ ≤ 1, “high impact, protein-quality” = Δ°K_D_ > 1 & ΔΔ°K_D_ ≤ 1, “high impact, protein-interaction” = Δ°K_D_ > 1 & ΔΔ°K_D_ > 1. C) DMS heatmaps of the ΔΔ°K_D_ (the relative difference in the shift of the °K_D_ of the antigen mutant compared to the antigen WT between the primary VHH compared to the control VHH) of single structural epitope mutations (top). Boxplots quantifying the ΔΔ°K_D_ per epitope position (bottom). The center line depicts the median and the box edges the 25th and 75th percentiles. The whiskers show the largest values no more than 1.5* the interquartile range.

For all antigen mutants, i.e., those in the epitope of the primary VHH, the change in °K_D_ of the primary VHH (Δ°K_D_) ranged between -0.68 (improved affinity) and 3.36 (decreased affinity) (Fig. 2A, B). 19%-52% of single mutations and 43%-74% of double mutations had a significant impact on the primary VHH with |Δ°K_D_| > 1. Of note, the main control VHH binding was impacted to an almost equal extend, with its Δ°K_D_ ranging from -0.64 to 3.75 and 14.6%-46.1% of mutations having a significant impact (|Δ°K_D_| > 1).

As mentioned above, each control VHH has a non-overlapping epitope to the primary VHH. Therefore, the control VHH Δ°K_D_ cannot be caused by *protein-interface* changes but must be a consequence of either allosteric effects, or *protein-quality* changes in the antigen. However, a single control VHH cannot be used to distinguish *protein-quality* changes from allosteric effects. Although allosteric effects have been shown to be abundant, they do not alter the entire protein equally ^24, 34^.

In contrast, *protein-quality* changes, which are equally prevalent ^28^, affect the entire protein. Therefore, to distinguish between these two mechanisms, all additional control VHHs with unique, non-overlapping epitopes were compared to the main control VHH (n_additional=2,1,3 for P0DPI0-7m1h_C, P0DPI0-7m1h_G and P0DTC2-7z1b_C respectively, Table 1). For P0DTC2-7olz_B only the main control VHH, but no additional controls were available. The average Spearman correlation was 0.96 (0.04 std) for all 6 additional controls to its corresponding “main” control, indicating that most antigen mutations measured here affect *protein-quality*, as opposed to allosteric effects or other local instability/unfolding events that don’t alter total protein abundance (Supp. Fig. 2A). Therefore, the main control Δ°K_D_ will from here on be used as a proxy for *protein-quality* changes.

In contrast to the control VHH, the primary VHH Δ°K_D_ consists of both *protein-quality* and *protein-interaction* changes. Therefore, when the primary and the control VHH Δ°K_D_ of an antigen mutation are highly similar (relative difference <= 1), the mutation alters only the antigen *protein-quality* (Fig. 2B). In contrast, any shift (more than 1) in the observed affinity of the primary VHH compared to the control VHH (ΔΔ°K_D_) reflects *protein-interaction* changes (Fig. 2B).

In the four studied complexes, ΔΔ°K_D_ ranged between 0 and 3.3 (>1000-fold-change in the primary VHH °K_D_ compared to the control VHH °K_D_, Supp. Fig. 1E). 43%-82% of the single mutations an 42-83% of double mutations that had an impact on the primary VHH (Δ°K_D_ > 1) had a ΔΔ°K_D_ of less than 1. This means that the Δ°K_D_ of the primary VHH and control VHH differ less than 10-fold, therefore suggesting that there was relatively little *protein-interaction* change caused by these mutants (Fig. 2B, 3A).

When aggregating mutations by position, 73%-93% of the primary VHH epitope positions exhibited comparable mutational constraint for both VHHs, i.e., >50% of singles had ΔΔ°K_D_ <1 (Supp. Fig. 4C). Conversely, 5%-27% of antigen positions had substantial impact on *protein-interaction* (>50% of singles had ΔΔ°K_D_ >1). Note that changes in *protein-quality* and *protein-interaction* are on a continuous scale and a mutation can influence both properties simultaneously, to different extents.

**Figure 3.**
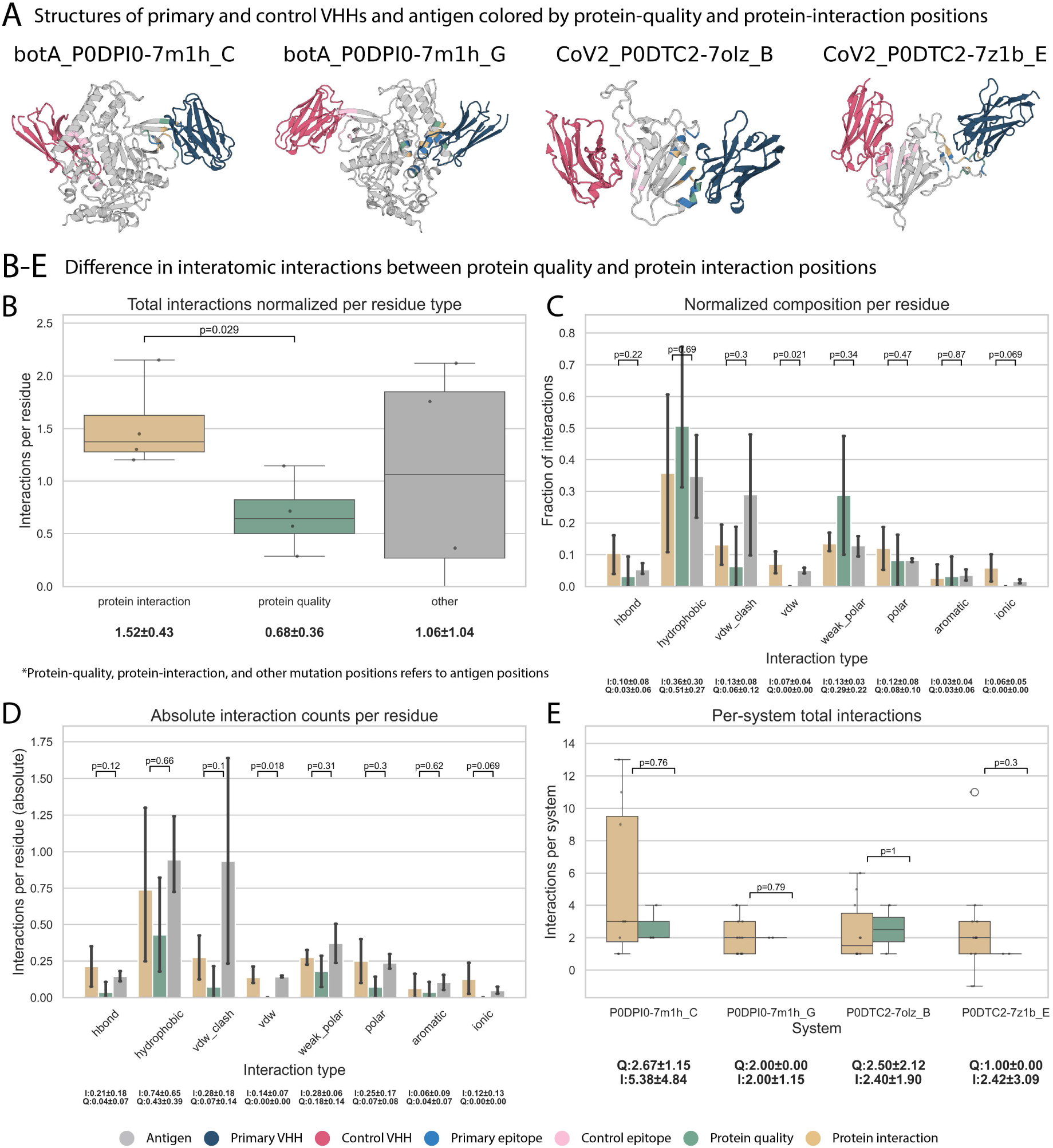
Protein-interaction mutations are characterized by charge changes and presence of bonds with the paratope residues. A) Complex PDB structures, colored by antigen, primary VHH, and control VHH. The mutated epitope positions are colored by their classification as “protein-quality”, “protein-interaction” or neither. B) Per complex, the mean normalized per residue group number of interactions over epitope positions of different classifications with the paratope. C) Per type of epitope position, the average fraction of interactions over the four complexes of each type of bond/interaction. D) Similar to C but showing the average absolute number of interactions instead of fractions. E) The total number of epitope-paratope interactions per complex.

**Figure 4.**
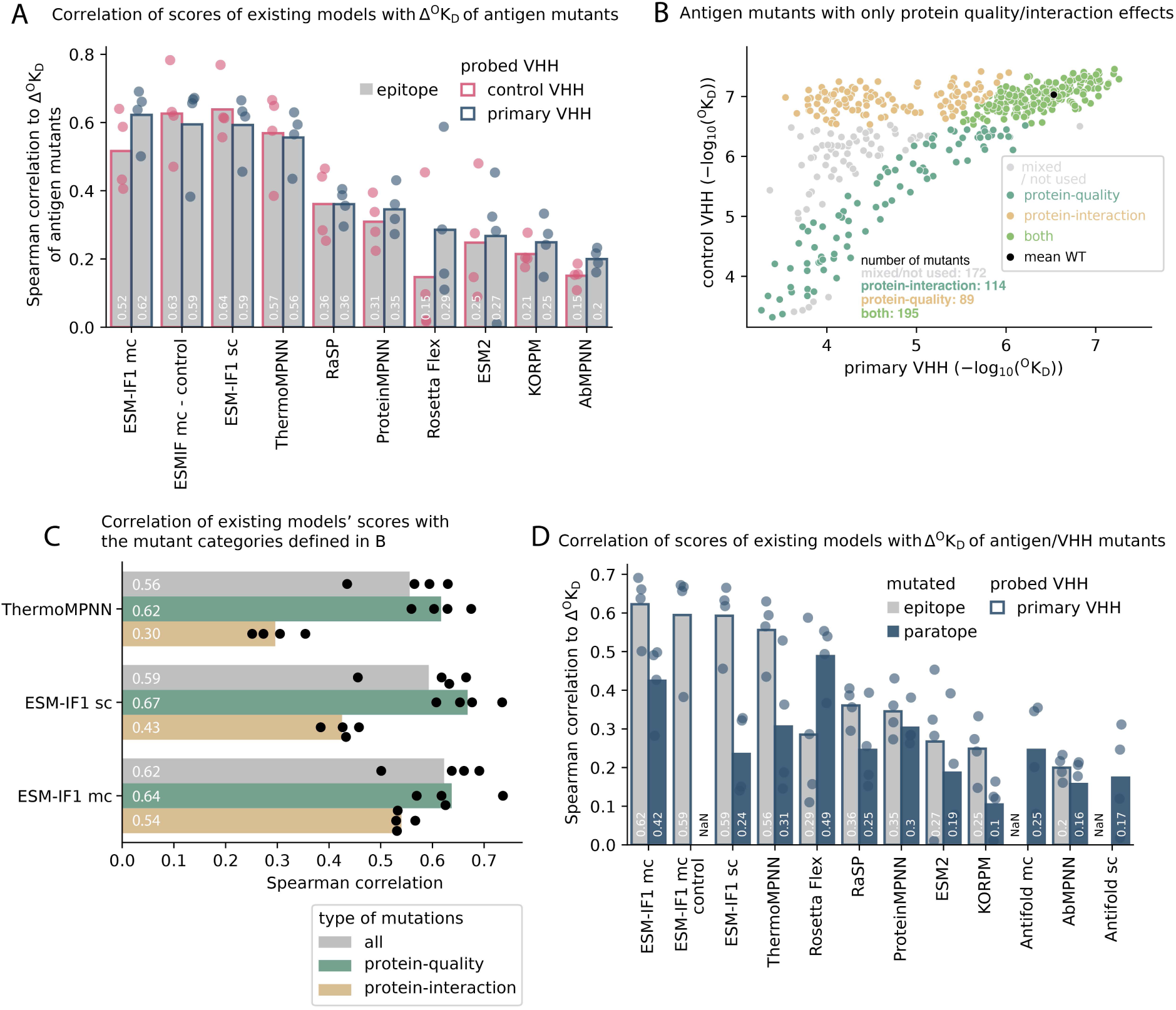
Existing models largely predict protein-quality not protein-interaction changes. A) The Spearman correlation between several models’ prediction scores to primary VHH and control VHH Δ°K_D_ of single structural epitope mutants. Each point is the correlation of the prediction score on all mutations of an antigen in a single complex from a given model. The bar indicates the mean of the four points. B) An example plot showing the classification of mutations based on primary VHH and control VHH Δ°K_D_ used to evaluate model performance on predicting changes in protein quality or protein interaction. Mutations with label “both” are used for either category. C) The Spearman correlations of ESM-IF1 and ThermoMPNN for all single epitope mutations, protein-quality epitope mutations, or protein-interaction epitope mutations (shown in B). D) Spearman correlation between several models’ prediction scores and antigen mutants (same data as show in the grey bars in A) and primary VHH mutants. Each point is the correlation between one model’s prediction score of all epitope mutations in a single complex. The bars indicate the mean of the four points.

To validate the high prevalence of *protein-quality* changes in observed affinity in non-VHH-antigen systems, interferon alpha (IFNA) bound by a Sifalimumab (IgG, PDB 4YPG) and the receptor IFNAR2 (PDB 3SE3) was investigated using AlphaSeq. IFNA was mutated, including the positions of IFNAR2 and Sifalimumab epitopes. Sifalimumab VH and VL were expressed as an scFv.

Next, the binding of these two proteins to all IFNA mutants was measured using AlphaSeq. The Δ°K_D_ of Sifalimumab ranged from -0.5 to 4.1 and for IFNAR2 from -1.0 to 3.9. The ΔΔ°K_D_ ranged from 3.2 (IFNAR2-specific) to -3.7 (Sifalimumab-specific). 86% of the mutations similarly affected the affinity of the two binders (ΔΔ°K_D_ <1), while 8% and 5% affected the binding of only IFNAR2 and Sifalimumab, respectively (*protein-interaction* mutations, ΔΔ°K_D_ >1/<-1), which matched the corresponding epitopes (Supp. Fig. 2B).

Taken together, similar to the epitopes of the four VHHs investigated above, only a small fraction of the epitope mutations of the Sifalimumab scFv and receptor IFNAR2 result in *protein-interaction* changes.

### Structural and biophysical analyses differentiate *protein-quality* and *protein-interaction* residue positions

To relate mutational effects to structural context, we first identified antigen positions showing the strongest separation between general effects on protein quality and interaction-specific effects (Fig. 2C). Positions whose substitutions similarly affected binding to both the primary and control VHHs were classified as *protein-quality* positions (lowest 20% of median ΔΔ°K_D_ across single mutants). In contrast, positions whose substitutions preferentially altered binding to the primary VHH were classified as *protein-interaction* positions (highest 20% of median ΔΔ°K_D_ across single mutants). All remaining positions were left unclassified.

To place these effects in a structural context, we visualized the complexes of the primary and control VHHs bound to their cognate antigens (Fig. 3A). Both *protein-quality* and *protein-interaction* positions were distributed across rigid secondary structure elements (β-strands and α-helices) as well as flexible loop regions of the epitopes, indicating that the observed mutational behavior cannot be explained solely by local backbone flexibility.

Hence, to characterize the biochemical and structural differences between these two classes, we examined physicochemical property changes associated with single amino acid substitutions and quantified the interatomic interactions formed by residues at *protein-quality* and *protein-interaction* positions. We did not find a strong statistically significant difference that is preserved across all VHH-antigen complexes. Subtle trends show that substitutions at *protein-interaction* positions more often involved charge-altering mutations, and, to a lesser extent, changes in polarity and residue size (Supp. Fig. 3A). Although the magnitude and direction of these effects varied across VHH-antigen complexes, substitutions at *protein-interaction* positions frequently involved smaller or more hydrophilic residues. In contrast, substitutions at *protein-quality* positions were dominated by property-preserving changes (conservative changes that preserve key biochemical properties) and showed little coordinated bias across biochemical features.

To further characterize these trends, we quantified the biochemical change introduced by each single amino acid substitution using continuous property differences (charge_diff, hydropathy_diff, and volume_diff; Methods). These metrics measure the magnitude of change in charge, hydropathy, or side-chain volume between the wild-type and substituted residue, allowing conservative substitutions to be distinguished from mutations that introduce larger biochemical changes. Unlike the categorical property analysis described above, these metrics capture subtle biochemical differences even when substitutions fall within the same property class (Methods, Table 4).

**Table 2:**
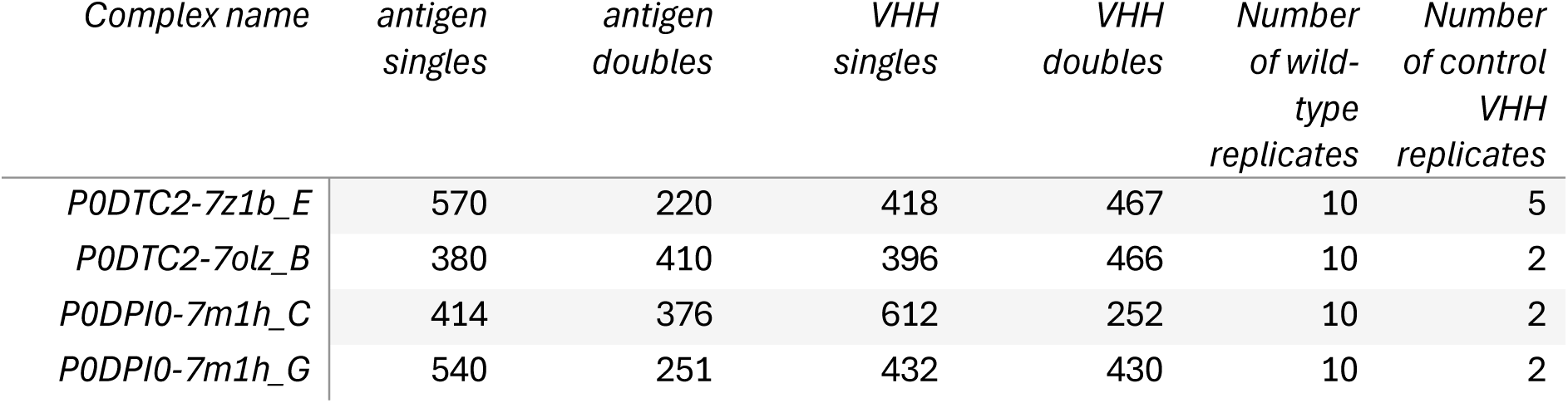
Number of single and double mutations in the antigen and VHH mutant libraries.

| Complex name | antigen<br>singles | antigen<br>doubles | VHH<br>singles | VHH<br>doubles | Number<br>of wild-type<br>replicates | Number<br>of control<br>VHH<br>replicates |
| --- | --- | --- | --- | --- | --- | --- |
| P0DTC2-7z1b_E | 570 | 220 | 418 | 467 | 10 | 5 |
| P0DTC2-7olz_B | 380 | 410 | 396 | 466 | 10 | 2 |
| P0DPI0-7m1h_C | 414 | 376 | 612 | 252 | 10 | 2 |
| P0DPI0-7m1h_G | 540 | 251 | 432 | 430 | 10 | 2 |

**Table 3:**
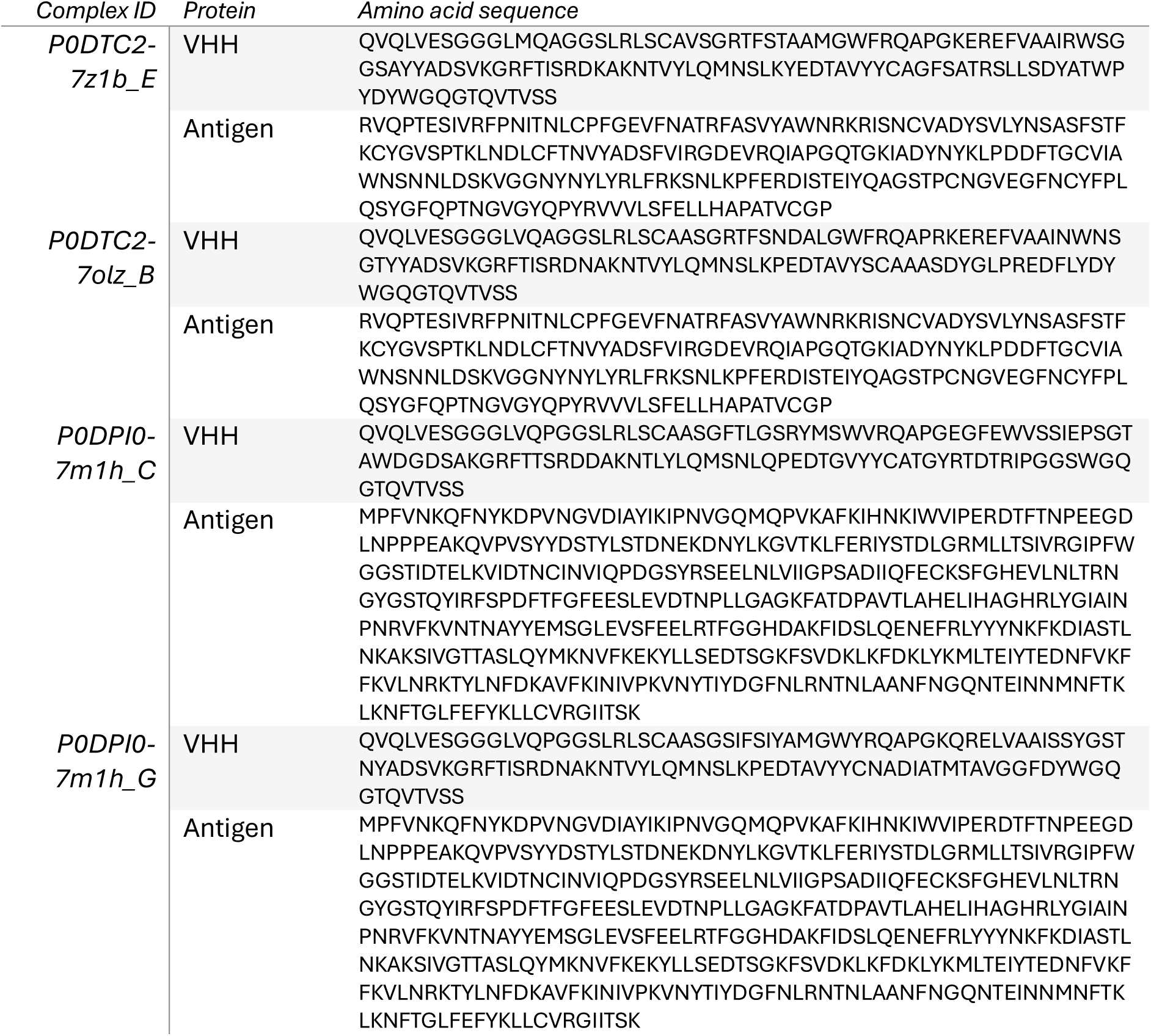
The expressed wild-type antigen and VHH amino acid sequences on yeast.

**Table 4:**
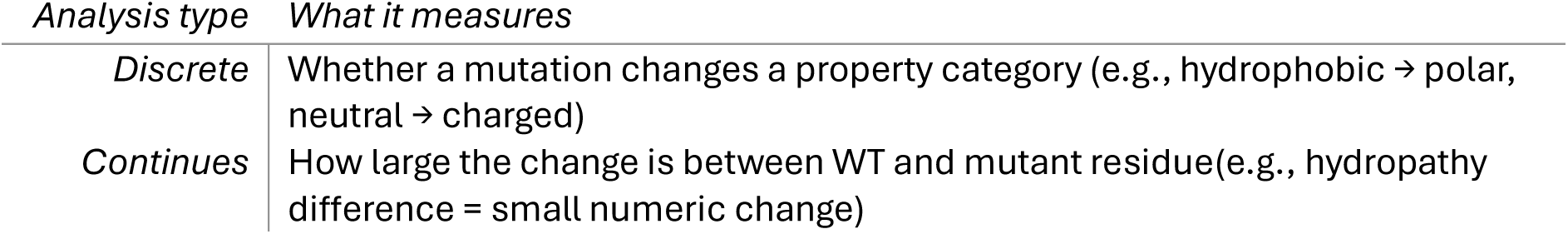
The difference between discrete and continues biochemical properties.

Statistical metrics (Methods) confirms that *protein-interaction* effects show modest but measurable deviations in charge and side-chain volume across several complexes, with marginal Mann-Whitney effects (average p ∼10^-2^, p-values ranging from 10^-1^ to 10^-5^) in P0DPI0–7m1h_G and P0DTC2–7olz_B, and a stronger hydropathy signal only in P0DPI0–7m1h_C (p = 2.3×10^-4^). Categorical analyses aligned with these continuous trends, showing system-specific enrichments for hydrophilic substitutions, smaller residues, and gains in positive charge or polarity (χ² p values ranging from 10^-2^ to 10^-7^). Despite these isolated signals, *protein-quality* substitutions are dominated by property-preserving changes, showing little evidence of systematic biochemical changes across substitutions. When aggregated across datasets, charge change remained the only globally significant feature, whether quantified as a continuous shift (p = 3.7×10^-3^) or as a categorical enrichment (χ² p = 1.4×10^-2^). Together, these results indicate that differences in how mutations affect °K_D_ across VHH-antigen pairings is most consistently driven by electrostatic or polarity-related substitutions, whereas hydropathy and volume effects, though detectable in specific complexes, exert more modest and context-dependent influences.

Structural mapping and interaction analyses revealed that *protein-quality* positions typically formed few or no direct interfacial bonds, consistent with mutations that proportionally affect binding to both primary and control VHHs (Fig. 3B, Supp. Fig. 3B). In contrast, *protein-interaction* positions exhibited significantly higher interaction densities, with pronounced enrichment in hydrogen bonds, polar, and ionic contacts, consistent with previous reports ^35^ (Fig. 3B-E). Specifically, *protein-interaction* positions were enriched in hydrogen bonds, polar, and ionic contacts, whereas *protein-quality* positions showed a relative enrichment in hydrophobic, aromatic, and weak polar interactions (Fig. 3C). Absolute interaction counts per residue mirrored these trends, with *protein-interaction* positions forming more contacts across most interaction types (Fig. 3D). Complexes containing a higher abundance of *protein-interaction* positions also displayed increased total interatomic interaction counts at the complex level, linking these positions to denser and more chemically specific interfaces (Fig. 3E).

By comparison, substitutions at *protein-quality* positions were predominantly conservative, preserving properties such as hydrophobicity, polarity, or side-chain size. Changes that alter these properties are more likely to affect antigen folding or stability, explaining the observed *protein-quality* effects. These results indicate that charge- and polarity-mediated contacts primarily drive *protein-interaction* effects, whereas hydrophobicity-based contacts underlie *protein-quality* behavior. Importantly, without control VHHs, affinity changes cannot be disentangled from global effects on antigen quality, making it impossible to confidently identify residues that directly govern epitope–paratope recognition.

### Inverse folding models primarily capture *protein-quality* changes of antigen mutants rather than *protein-interaction* changes

There exist a variety of models that can score the effect of protein mutants as ΔΔG, likelihoods, or similar scores. To evaluate if these models can disentangle *protein-quality* changes and *protein-interaction* changes that underlie the primary VHH Δ°K_D_, we used the AlphaSeq data presented here to benchmark a set of models with different architectures. In this benchmark, we included structure-conditioned inverse-folding models (ESM-IF1, ProteinMPNN, AbMPNN) ^7, 36, 37^, models focused on predicting ΔΔG_stability_ (RaSP, KORPM, ThermoMPNN) ^38–40^, *Rosetta Flex* ^41^, which predicts ΔΔG_interaction_, and a sequence-only protein language model (pLM, ESM2) ^42^. We compared each model’s score, e.g., pseudo-perplexity, log-likelihoods, or interface/stability (ΔΔG) prediction scores, with experimentally measured affinities of single-mutant antigen variants in complex with the respective wildtype VHH.

ThermoMPNN and ESM-IF1 had the highest (anti)correlation to primary VHH Δ°K_D_ (Spearman’s R=-0.56 for ThermoMPNN, and 0.59/0.62 for ESM-IF1 single-chain (sc) / multi-chain (mc) model, respectively, p=0.18/0.07 t-test ThermoMPNN vs ESM-IF1 sc/mc, Fig. 4A, Supp. Fig. 5C). Indeed, ESM-IF1 log-likelihood anticorrelated with ThermoMPNN (Spearman’s R=-0.64), with low likelihood corresponding to high predicted ΔΔG_stability_ (Supp. Fig. 5A). In addition, both underperformed on the epitope of P0DTC2-7z1b_E, one of the SARS-CoV-2 Spike RBD VHHs. These results suggest that these two models at least partly predict the same protein characteristics. ThermoMPNN/ESM-IF1 scores correlated better to Δ°K_D_ than Rosetta Flex’ predicted ΔΔG_interface_ for three out of four complexes (average Spearman’s R=0.29, Fig. 4A). This is in agreement with the finding based on ΔΔ°K_D_ that *protein-quality* changes generally drive changes in observed affinity of antigen mutants and suggests that ThermoMPNN and ESM-IF1 both primarily predict *protein-quality* changes.

The scores of the other two models that predict ΔΔG_stability_ had moderate correlation with ThermoMPNN and each other (Spearman’s r=0.44-0.71, Supp. Fig. 5A). In addition, the top 10% predictions of the three models overlapped substantially (45-53%, compared to 0-37% with other models, p=2.3*10^-5^ t-test), increasing the confidence that these models all predict the same protein characteristic, i.e., stability (Supp. Fig. 5B). Still, ThermoMPNN’s score had a significantly better anticorrelation to Δ°K_D_ than the other two models (Spearman’s R of -0.36 for RaSP and -0.25 for KORPM, p=0.034 t-test, Fig. 4A, Supp. Fig. 5C).

If a model can predict *protein-interaction* on top of *protein-quality*, it should perform better on the primary VHH compared to the control VHH, which binds distal to the mutated sites and which interface is not directly affected by the mutation. Therefore, we compared the models’ performance on the effect of antigen mutants on the primary VHH and control VHH Δ°K_D_. Most models, including ESM-IF1 single-chain mode predicted the control VHH Δ°K_D_ as well as the primary VHH Δ°K_D_ (Spearman’s R=0.64, 0.59 respectively, Fig. 4A) supporting the hypothesis that for ESM-IF1, similarly to ThermoMPNN, the predictive signal of antigen mutants mostly results from predicting changes in *protein quality*, not *protein interaction*. The high performance of ThermoMPNN on predicting the control VHH Δ°K_D_ / *protein-quality* is in line with previous literature that shows that protein expression on the yeast surface (i.e. *protein-quality*) is correlated to the thermostability of a protein^43^.

The fact that ESM-IF1 primarily predicts *protein-quality* was confirmed for non-VHH protein-protein interactions on the previously introduced dataset of IFNAR2 receptor (primary) and Sifalimumab (control) binding IFNA. The single mutants of the IFNAR2 epitope positions were scored by ESM-IF sc conditioned on the antigen chain of the IFNAR2 complex structure. ESM-IF1 log-likelihood correlated almost equally to the Δ°K_D_ of both binders, the “primary” IFNAR2 and the “control” Sifalimumab (0.74 vs 0.70 Spearman’s correlation). ThermoMPNN had similar correlations to Δ°K_D_ under the same conditions (−0.75 vs -0.66, Supp. Fig. 5F).

Notably, the choice of experimental X-ray structure strongly affected the performance of both models. For Sifalimumab epitope mutations and Sifalimumab Δ°K_D_, ESM-IF1 prediction improved when it was conditioned on the antigen chain from the IFNAR2 structure rather than from the antigen chain from the Sifalimumab structure itself (Spearman’s R: 0.70 vs 0.51 for ESM-IF1; -0.66 vs -0.5 for ThermoMPNN).

This was confirmed using a previously published AlphaSeq dataset of affinities of 33 SARS-CoV-2 binding IgGs/VHHs to all possible single SARS-CoV-2 mutants^31^. The 24 antibodies for which an antibody-antigen structure was available were analyzed. ESM-IF1 sc log-likelihood (LL) was calculated 24 times for all single RBD mutants, each time conditioning on the antigen chain from a different antibody-antigen structure. For each antibody, the Δ°K_D_ of mutations in its epitope was correlated to each of the 24 ESM-IF1 prediction sets. Only 1 of the 24 antibodies had the highest correlation of LL to the Δ°K_D_ when ESM-IF1 was conditioned on the antigen chain of its own structure. For the other 23 antibodies, the highest correlation was achieved using a non-matching antigen structure. Across these antibodies, only six unique antigen structures yielded the highest correlation. These six structures did not have significantly better resolution than the other 18 structures, nor were the structures more similar to the apo (unbound) conformation.

When analyzing not only the highest, but all Spearman correlations of 24 sets of ESM-IF1 LL and Δ°K_D_, the mean and spread varied widely per antibody (Supp. Fig. 5G). This variability could stem from differences in the biophysical complexity of the associated interface, or from differences in the closest data point(s) in the training data of ESM-IF1.

So, even though the conformation of the RBD might be slightly different in the holo (bound) state per antibody, other, yet to be determined, characteristics of the experimental structure appears more important for ESM-IF1 prediction accuracy.

The results above suggest that existing models are better at predicting *protein-quality* changes than *protein-interaction* changes. To quantify this directly, predictive scores of the models were separately correlated to the Δ°K_D_ of mutants with only *protein-quality* changes, or mutations with only *protein-interaction* changes (Fig. 4B). The predicted ΔΔG_interaction_ from Rosetta Flex was lower for the first group, and unchanged for the second group, confirming that these mutants indeed represented changes in *protein-quality* and *protein-interaction* (Fig. 4C). A non-significant increase in performance was observed for ThermoMPNN and ESM-IF1 single-chain for *protein-quality mutants* compared to all mutants (p=0.20/0.28 t-test, Fig. 4C). For *protein-interaction mutants,* these two tools had consistently lower (anti-)correlation with Δ°K_D_ (p=0.005, Fig. 4C). Notably, the drop in performance for *protein-interaction* mutations was less pronounced for ESM-IF1 multi-chain mode (p=0.086 t-test), which outperformed all models including ThermoMPNN on this subgroup (Fig. 4C).

When two mutations are introduced in a protein simultaneously, one mutation can influence the effect of the other mutation on protein function, e.g., protein binding. This effect, where the two mutations are “non-additive”, i.e., their combined change in free energy (ΔΔG, directly derived from and related to the Δ°K_D_, see Methods) is not simply an addition of the ΔΔG of the two single mutations, is known as *epistasis* ^27^. This phenomenon is therefore essential for a model to capture if it is applied for affinity maturation or *de novo* design. Therefore, we use double mutations to further benchmark the capabilities of ESM-IF1, the model with the best overall ability to predict Δ°K_D_, which can also predict the effect of multi-site mutations. The correlation of ESM-IF1 log-likelihoods with Δ°K_D_ was lower for double compared to single mutants (r = 0.44 vs 0.59, p=0.03 t-test, Supp. Fig. 5D). In addition, we investigated the performance on predicting ΔΔG of additive doubles (where the ΔΔG is the same as the sum of the ΔΔG of the two singles) and non-additive doubles. ESM-IF1 performed better for additive compared to non-additive mutants (r=0.43 vs 0.18, p=0.02 t-test, Supp. Fig. 5E). Together, these results indicate that while ESM-IF1 is useful in ranking single-mutant variants for affinity maturation, its utility for higher-order mutants is likely substantially lower.

In conclusion, out of the evaluated models, ESM-IF1 has the best overall predictive power for observed affinity changes of antigen mutants. It performs on-par or slightly better than ThermoMPNN. The results indicate that ESM-IF1 performs best at predicting changes in *protein-quality* for single mutants but is more limited in its ability to capture *protein-interaction* effects or the impact of double mutants, particularly when epistatic interactions are involved.

### Inverse folding models fail to capture mutational effects in VHHs

Having examined the predictive performance of state-of-the art models on antigen mutations, we next evaluated their ability to predict the effects of mutations within the VHH. Importantly, VHH mutations Δ°K_D_ could not be separated into *protein-quality* and *protein-interaction* changes as no control binder that bound the VHHs outside of the paratope was available. However, mutations in the loops of the complementarity-determining regions (CDRs) of antibodies are likely to have less impact on *protein-quality* than mutations in secondary structures of antigens ^44^. As most of the mutations that we introduced in the VHHs are in the CDRs (97% of singles, and 100% of doubles), fewer unstable variants are expected.

Consistent with this expectation, the predicted ΔΔG_interaction_ from Rosetta Flex showed a higher correlation with the Δ°K_D_ of VHH mutants than antigen mutants for three out of four VHH-Ag complexes (p=0.13, Fig. 4D). In addition, the scores of all other tested models, including ESM-IF1, had lower correlations with the Δ°K_D_ of VHH mutants (Spearman’s R=0.09-0.31) than antigen mutants (R=0.15-0.59) (p=0.03, Fig. 4D, Supp. Fig. 6). ESM-IF1 multi-chain mode improved the correlation for VHH mutants by 18 percentage points to 0.42, which is a lower performance than ESM-IF1 single-chain mode for antigen mutants (Spearman’s R=0.24). In conclusion, existing models have less predictive power on the change in observed affinity of VHH mutants compared to antigen mutants.

Our results suggest that this is due to the larger effect of paratope mutations on *protein-interaction* and the lower impact on *protein-quality* compared to epitope mutations, and the inability of existing models to predict *protein-interaction* changes accurately.

## Discussion

Here, we present a large high-throughput mutational scanning yeast-display dataset of VHH-antigen affinities and use a control VHH to separate *protein-quality* from *protein-interaction* changes. We show that the majority of the measured signal reflects changes in *protein-quality* arising from mutations and the percentage of such mutations differs by complex. Our dataset covers four VHHs with distinct, non-overlapping interfaces and two different antigens, SARS-CoV-2 spike RBD, and Botulinum Toxin A.

*Protein-interaction* changes were mostly detected for mutations at positions in the interface that formed a bond with the binding partner. In contrast, mutations at interface residues that do not form bonds predominantly affect the *protein-quality* of the mutated protein. *Protein-quality* changes therefore dominate the observed affinity signal. Using the two components underlying observed affinity, we show that existing models ThermoMPNN and ESM-IF1 can already capture *protein-quality* effects with reasonable accuracy, indicating that sufficient data is available to have learned this protein property. However, they can only predict little of the *protein-interaction* changes in the dataset (Figure 4C). A number of factors could contribute to the inability of current models to predict context-dependent effects that govern antigen recognition. First, the structural priors used by inverse folding models are typically learned from static crystal structures, which poorly represent the flexibility and conformational diversity of CDR loops. Second, antibody sequences, particularly single-domain VHHs, are underrepresented in the training data of most models, limiting their generalizability. Third, inverse folding scores reflect compatibility with a fixed backbone, which do not take into account the side chains and thus the interface bonds, which are essential for VHH-antigen binding.

As existing models can already predict *protein-quality* changes with reasonable accuracy, collecting more data that is dominated by *protein-quality* effects is unlikely to substantially improve affinity prediction models. Instead, large scale affinity measurements should incorporate the correct controls or complementary stability/expression assays that disentangle *protein-quality* from true *protein-interaction* changes. Such data would enable the efficient training of the next generation of models that learn true protein-protein interaction, in addition to general protein-quality features.

The data that was presented here are from yeast-display mating assays that are influenced by *protein-quality*, i.e., by the number of correctly folded molecules. The influence of *protein-quality* on observed affinity will depend on the affinity assay being used. Public databases of protein mutation data, e.g., SKEMPI, mostly contain data of traditional methods of affinity measurements, like Surface Plasmon Resonance (SPR) readouts^29, 45^. Because these measurements are performed with purified proteins, the resulting affinities are largely independent of expression levels or the fraction of protein that is active. However, this correction might often not have been applied in ^23^. In addition, traditional biophysical methods cannot practically generate the thousands of datapoints needed for machine learning models, requiring high-throughput assays. These assays are typically display-based (yeast, phages, or ribosomes) and influenced by protein-quality, making it essential to measure correctly folded (active) protein alongside affinity within the same assay ^30^.

The next step in antibody affinity modeling is to use large-scale DMS datasets in which protein-interaction effects are separated from protein-quality effects to train new models, or fine-tune existing ones, to better learn patterns of binding. In principle, models trained on multiple binders or control VHHs could also learn to disentangle these effects computationally, but experimentally resolved datasets provide a clear foundation for such approaches.

### Limitations of this study

As universal anti-VHH binders are scarce, we were not able to add the appropriate controls to experimentally quantify the contribution of *protein-quality* changes to the Δ°K_D_ values of VHH mutants. Even so, predicted interface and stability ΔΔG values suggest that VHH variants are less affected by *protein-quality* than antigen Δ°K_D_, implying that VHH DMS datasets may be more suitable for model training, even in the absence of explicit controls. Such controls would, however, likely further improve the fidelity of the affinity signal.

For the analysis of biochemical features of *protein-interaction* and *protein-quality* positions, positions with the top and bottom 20% median ΔΔ°K_D_ values were used, while both are continuous features without true cut-offs.

## Methods

### Complex selection

Single-chain antigens were selected from SabDab^46^ for which one structure was available with multiple VHHs binding different epitopes of the same antigen (i.e., 7m1h) or multiple structures were available of the same antigen with each containing a VHH that bound a different epitope (i.e., 7olz, 7z1b). For each selected antigen, a primary VHH and one or multiple control VHHs were selected that had non-overlapping epitopes (see below). If more than one control VHH was selected, each had a distinct epitope. The “main” control VHH that was used for analyses was the control VHH of which the epitope had the largest geometric distance from the primary VHH epitope in the antigen structure of the primary VHH.

### Antigen and antibody sequences

For antigens, the canonical human amino acid sequence from UniProt was used as a reference. For VHHs, the sequence was taken from the complex’ SabDab *fasta* file, and ANARCI ^30, 46^ was used to confirm that each sequence only contained the VH (i.e. no tags etc).

### IFNA complex

In addition to the four VHH complexes, the Sifalimumab and IFNAR2 in complex with IFNA were also analyzed. 1536 single mutations in 81 positions of IFNA were introduced in the epitope of Sifalimumab and IFNAR2 and all residues that were positions in the amino acid sequence between these two sets of epitope residues. The affinity of all of these IFNA mutants were then measured for both Sifalimumab and IFNAR2 using AlphaSeq.

### VHH-antigen structure processing

The VHH and antigen chain of interest were extracted from the SabDab *pdb* file. Each chain was renumbered to have 1-indexed consecutive numbering. Using *PDBFixer v1.9*^47^, the structure was processed in the following steps: heterogens were removed, missing heavy atoms were added, missing hydrogens were added and non-standard residues were replaced *(functions: ’findMissingResidues’, ’findNonstandardResidues’, ’replaceNonstandardResidues’, ’removeHeterogens(False)’, ’findMissingAtoms’, ’addMissingAtoms’,’addMissingHydrogens(7.0)’)*. The renumbered structure was used for most steps and only for the biophysical analysis was the *PDBFixer*-processed structure used. For interatomic interaction calculations, intermediate PDB files were generated containing three chains (primary VHH, antigen, and control VHH) merged into a single structure. Chain identifiers were renamed as A for antigen, B for primary VHH, and C for control VHH to ensure consistency and facilitate downstream processing by custom analysis scripts.

### Epitope and CDR definitions, single and double mutations

The “strict” epitope was determined as any residue containing at least one atom within 4.5Å of any VHH atom. The “inclusive” epitope is defined as any residue containing at least one atom within 5Å of any VHH atom and/or a Cα atom within 10.5Å of any VHH atom. Identical distance-based criteria relative to the antigen were applied to define the paratope on the VHH. CDRs are assigned using ANARCI^46^ according to the IMGT^48^ CDR definition (CDRH1: 24-40, CDRH2: 55-66, CDRH3: 103-119). “Strict” and “inclusive” definitions of the epitope and paratope were created for single and double mutant selection only (see next paragraph).

### Mutant selection

A library of mutants was designed for both the paratope and the epitope. For each, all possible single mutants of the “inclusive” epitope definition were included in each library (Table 2). In addition, ten synonymous wild-type variants were added for the VHH and the antigen. Moreover, two to five synonymous variants of each selected control VHHs were added. Doubles were selected within the “strict” epitope definition to fill the capacity of the library (Table 2). Doubles were enriched for a number of metrics or model predictions:

For P0DTC2-7z1b_E: NanoBERT^49^ (VHH) or ESM2^42^ (antigen) perplexity, MM/GBSA^50^ predicted ΔΔG, alanine-scanning °K_D_ values, pair-wise geometric distance.

For the other complexes: ESM-IF1^7^ log likelihood instead of pLM perplexity, Rosetta Flex ΔΔG^51^ instead of MM/GBSA, ThermoMPNN^40^, RaSP^38^, and KORPM ΔΔG, alanine-scanning °K_D_ values, and pair-wise geometric distance.

All scores were scaled to 0-1, and a composite score 𝑐𝑠 was calculated using the geometric mean of all individual scores. The library was generated by sampling new mutations one-by-one with weight 𝑐𝑠^𝑖^, where 𝑖 = 5 for P0DTC2-7z1b, and 𝑖 = 3 for the other complexes.

The chance of the sampled mutation to the library was calculated as:

Let the considered double mutation 𝑀 consists of single mutations 𝑚_1_, 𝑚_2_ with corresponding wild-type residues 𝑤_1_, 𝑤_2_, and 𝐷(𝑎, 𝑏) be the BLOSUM62 distance between residues 𝑎 and 𝑏. For each subject mutation 𝑖, compute:

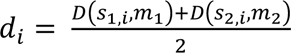 and 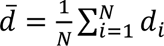 for 𝑁 previously sampled mutations, and finally 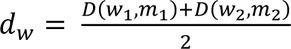, then similarity score 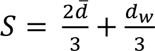, and sampling probability 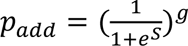 with 𝑖 = 1 for P0DTC2-7z1b, and 𝑖 = 5 for the other complexes.

This approach aimed to simultaneously diversify the amino acids that were selected using the BLOSUM62 distance, and to decrease the number of non-binder doubles using the other metrics and model predictions. 𝑖 and 𝑔 were updated after evaluation of the data of P0DTC2-7z1b, and before generating the libraries for the other systems.

### Yeast-display affinity measurement

The A-Alpha Bio assay “AlphaSeq” was used to calculate °K_D_ values using the library of single and double mutants for the epitope and paratope, as described previously (<u>Younger</u>, <u>Engelhart</u>, <u>Engelhart</u>). Expression was measured by binned FACS sorting and NGS as described previously. Observed affinity was reported as °K_D_ = − log_10_ 𝐾_𝐷_ where 9 corresponds to 1nM and 6 to 1μM. The wild-type sequences expressed in this assay can be found in Table 3.

### Affinity measurement processing

AlphaSeq measures the °K_D_ of all VHH mutants with all antigen mutants, creating an all-vs-all matrix of °K_D_ measurements. Here, we extract the data columns from the AlphaSeq library-on-library data of WT primary or control VHH binding antigen mutants, and rows of VHH mutants binding WT antigen. In addition, we used WT-WT binding measurements.

Although expression varied little (Supp. Fig. 7A), to ensure consistency of the data, any mutant with a measured expression of mean-bin lower then 1.75 were removed from any analyses, except for the Δ°K_D_ heatmaps in Fig. 2/Supp. Fig. 1.

For each antigen mutant, the °K_D_ was measured ten times, once for each synonymous variant of the wild-type primary VHH. Therefore, for all analyses, these ten measurements were mean-pooled. Mean-pooling was similarly applied for the two to five synonymous variants of each control VHH and each antigen mutant (separately for each control VHH), and lastly for the primary VHH mutants over the 10 synonymous variants of the antigen wild-type. The wild-type °K_D_ for each complex was calculated as the average of the 100 wild-type VHH wild-type antigen °K_D_ measurements (each of the 10 synonymous wild-type VHH variants binding each of the 10 synonymous wild-type antigen variants).

For each epitope and paratope mutant, a Δ°K_D_ was calculated as Δ°K_D_ = °K_D,mutant_ − °K_D,wild−type_, thereby separating mutation-induced shifts in affinity from intrinsic complex-to-complex variation. A negative Δ °K_D_ reflects a mutant protein that has a worse affinity for the binding partner. As Δ °K_D_ is measured in log_10_ space, a Δ°K_D_ of -1 reflects a decrease in affinity by a factor of 10.

Furthermore, a ΔΔ°K_D_ was calculated as: ΔΔ°K_D_ = | Δ°K_D_ target - Δ°K_D_ control |. Mutants where ΔΔ°K_D_ == 0 exhibit an equal change in observed affinity in both the control and primary VHH and therefore predominantly reflecting changes in *protein-quality* of the mutated protein. Any ΔΔ°K_D_ away from 0 reflects the extent to which a mutation differentially perturbs the primary VHH binding versus the control VHH binding, and thus serves as a proxy for epitope-specificity. Variants were ranked by their median ΔΔ°K_D_ across replicates, with those in the lowest 20th percentile classified as *protein-quality mutants* and those in the highest 20th percentile classified as *protein-interaction mutants*. Variants with intermediate deviations were not assigned to either category.

To calculate the performance of models on *protein-interaction* vs *protein-quality* the model prediction scores could not just be correlated to the Δ°K_D_ of the mutants of each group, as the distribution of the underlying °K_D_ values was different for each category for most complexes. This is problematic, as the standard deviation of °K_D,_ differs over the °K_D_ range, as measured for each mutant of the 10 synonymous wild-type variants of the binding partner (Supp. Fig. 7B). To resolve this, we selected variants that were max 0.5 away from a line of slope 1 where x=°K_D,primary_ and y=Δ°K_D,control_ and where the line passes through the point of the wild-type antigen. For these mutants, the protein interaction is assumed to be not affected or to a minor extent and are therefore labeled *protein-quality.* Similarly, a line is drawn through the antigen-wildtype where the Δ°K_D,control_==0 and mutants less than 0.5 away from this line (in Δ°K_D,primary_ space) are selected as *protein-interaction mutants.* For each group, the primary VHH Δ°K_D_ values of these mutants are correlated to a tool’s prediction score. For both groups, the range of °K_D,primary_ and, therefore, noise level distributions are similar.

Additivity was evaluated based on ΔΔG of single and double mutations. ΔΔG = 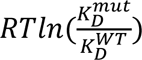 with the gas constant 𝑅 = 8.314 𝐽 𝐾^−1^𝑚𝑜𝑙^−1^, and the temperature taken as 𝑇 = 298. As the change in K_D_ is defined as 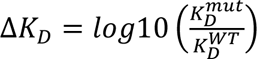, this means that ΔΔG = 1.364 Δ𝐾_𝐷_. Non-additive doubles of *mut1,mut2* were defined as having ΔΔ𝐺_𝑚𝑢𝑡1,𝑚𝑢𝑡2_ > 1 + ΔΔ𝐺_𝑚𝑢𝑡1_ + ΔΔ𝐺_𝑚𝑢𝑡2_.

### Public IgG/SARS-CoV-2 AlphaSeq dataset processing

In addition to the AlphaSeq data processed as described above, a previously published AlphaSeq dataset of VHHs and IgG binding SARS-CoV-2 spike RBD^31^ was re-analyzed. The previously processed dataset contains the 33 antibodies and 2434 single mutant variants on 165 positions of the RBD.

For 24 antibodies, a matching PDB structure was identified with an antibody-antigen complex. The epitope of each antibody was determined by taking the residues within 5 angstrom of the antibody chain(s). Each epitope was then mapped onto the high-resolution RBD structure PDB 7K9Z, chain E. 6 non-overlapping “primary” epitopes were selected. For each of these, a set of control antibodies was selected that did not overlap with the “primary” epitope itself, and that had an °K_D_ of 7 (100 nM) or stronger. Between 2 and 9 controls were identified. For each primary epitope, the mutations of that epitope’s positions were then used to correlate the °K_D_ of all controls amongst each other.

### Model evaluation

The following models were run to for the single/double mutants of the antigen/VHH:

- ThermoMPNN^40^ (https://github.com/Kuhlman-Lab/ThermoMPNN, September 2025, single mutations only)
- KORPM^39^ (https://github.com/chaconlab/korpm, September 2025, single mutations only)
- RaSP^38^ (https://github.com/KULL-Centre/_2022_ML-ddG-Blaabjerg/, September 2025, single mutations only)
- ESM-IF1^38^ (fair-esm v2.0.0, model: esm_if1_gvp4_t16_142M_UR50, scoring function: ‘score_log_likelihoods.py’)
- AntiFold^52^ (https://github.com/oxpig/AntiFold, September 2025, VHH mutations only)
- ProteinMPNN^36^ (https://github.com/dauparas/ProteinMPNN, September 2025, model: v_48_020)
- AbMPNN^37^ (code from ProteinMPNN, weights from https://zenodo.org/records/8164693)
- ESM2^42^ (https://github.com/facebookresearch/esm)^42^
- Rosetta Flex^51^ (https://github.com/Kortemme-Lab/flex_ddG_tutorial, with settings “nstruct: 15, backrubntrials: 5000, backrubtrajstride: 1000”)

ThermoMPNN, KORPM, RaSP, ESM-IF1, AbMPNN were all run with default settings. AntiFold log-likelihoods were generated in an autoregressive manner using the <u>score_log_likelihoods.py</u> function from ESM-IF1 and the model weights from AntiFold.

In addition, ESM-IF1 single-chain log-likelihoods and ThermoMPNN ΔΔG were calculated for all IFNA mutants, and correlated to the °K_D_ of either IFNA binder.

Finally, for the IgG/SARS-CoV-2 dataset, ESM-IF1 single-chain was run using default settings on all RBD mutants, and for each of the SARS-CoV-2 spike RBD chains of the 24 structures. This resulted in 24 sets of ESM-IF1 log-likelihoods. For each of the 24 antibodies, the mutants of that antibody’s epitope were used to correlate the °K_D_ of that antibody to the ESM-IF1 log-likelhoods conditioned on the RBD chain of that antibody’s antibody- RBD complex structure. This was then repeated 23 times, each time using a set of EMS-IF1 log-likelihoods conditioned on the RBD chain of one of the other antibody-antigen complex structures.

### Determination of protein-quality and protein-interaction positions

Positions were ranked by their median ΔΔ°K_D_ across single mutations at each position, with those in the lowest 20th percentile classified as *protein-quality* (global effects) and those in the highest 20th percentile classified as *protein-interaction* (epitope-specific effects). Variants with intermediate deviations were not assigned to either category.

For single-mutant entries, mutation type strings were parsed to extract residue position, wild-type residue, and mutant residue using regular-expression–based parsing. These annotations were used to generate position-resolved visualizations, including scatterplots, heatmaps, and boxplots.

### Biophysical properties

Each single amino acid mutation ID string was parsed using a regular expression, extracting wild-type residue name, position, and mutant residue name. For each mutation, we derived five high-level biochemical descriptors: amino-acid similarity class, polarity change, charge change, volume change, and hydropathy change, using Kyte-Doolittle hydropathy scores.

First, the amino-acid similarity class was determined by assigning each residue to one of four biochemical groups: non-polar/hydrophobic (G, A, V, L, I, M, P, F, W), polar uncharged (S, T, C, Y, N, Q), acidic (D, E), and basic (K, R, H). A substitution was considered conservative when the wild-type and mutant residues belonged to the same group, reflecting preservation of similar chemical properties (e.g., L-->I within the hydrophobic class or E-->D within the acidic class). A mutation was labeled semi-conservative when the residues belonged to different but chemically related groups, specifically transitions within the charged classes (acidic <--> basic) or within uncharged classes (non-polar <--> polar). These substitutions preserve partial biochemical similarity while altering certain functional attributes. All remaining substitutions (those crossing between unrelated groups, such as hydrophobic to acidic or aromatic to basic) were classified as radical, indicating maximal biochemical departure. Any substitutions involving residues not represented in the predefined groups were also treated as radical.

Second, polarity change was assessed by defining residues as either polar (all polar, acidic, and basic residues) or non-polar. A substitution was classified as having gained polarity if the mutant residue was polar while the wild-type was non-polar, and as having lost polarity in the opposite case. Mutations in which both residues shared the same polarity classification were marked as having no polarity change. This descriptor reflects alterations in hydrogen-bonding capacity and solvation.

Third, charge change was quantified using a discrete side-chain charge scale in which K and R were assigned a charge of +1, D and E a charge of -1, and all other residues (including H, due to its context-dependent protonation state) were treated as neutral. The net charge difference (mutant minus wild-type) was used to categorize each substitution as becoming more positive, more negative, or unchanged, thereby capturing potential modifications to local electrostatics.

Fourth, volume change was computed using IMGT side-chain van der Waals volumes. Mutations were labeled as larger or smaller depending on whether the mutant residue’s volume exceeded or fell below that of the wild-type residue. Identical volumes or unresolvable cases were treated as having no change. This metric highlights steric alterations that may impact packing, structural stability, or interface complementarity.

Finally, hydropathy change was computed from Kyte-Doolittle hydropathy scores. The hydropathy difference between mutant and wild-type residues determined whether the substitution became more hydrophobic, more hydrophilic, or remained unchanged. This descriptor captures alterations in hydrophobic character that influence folding energetics, solvent exposure, and antigen-antibody binding behavior.

For each feature, a categorical label (Table 4) was assigned, representing the directionality of the biochemical change. Numeric encodings were additionally computed as the difference between mutant and wild-type charge (charge_diff = charge_mut - charge_wt), hydropathy (hydropathy_diff = hydropathy_mut - hydropathy_wt), and side-chain volume (volume_diff = volume_mut - volume_wt).

For numeric (Table 4) biochemical feature distributions (charge_diff, hydropathy_diff, volume_diff) violin plots were used to compare charge_diff, volume_diff, and hydropathy_diff between *protein-quality* and *protein-interaction* mutants. Mann-Whitney U tests were used to assess differences between groups. For categorical biochemical distributions (charge_type, polarity_change, charge_change, size_change, hydropathy_change), we computed per-class proportions for each descriptor along with Wilson 95% confidence intervals. Bars were annotated with mutation counts. A global χ² test was used to compare distributions between diagonal classes, and per-category enrichment was separately tested using χ² or Fisher’s exact test depending on expected cell counts.

### Interatomic interactions

For each antigen-antibody PDB entries, we gathered the final primary VHH-antigen-control VHH complex by aligning separate entries by the antigen. Structural interaction data were derived from the merged complexes by annotating residues as paratope and epitope based on 4.5 angstrom distance between each antibody and antigen in the respective complex. Residues were grouped into two antigen epitope regions (primary epitope and control epitope) and two antibody paratope regions corresponding to primary and control VHH. In addition, each complex included curated sets of *protein-*quality and *protein-interaction* mutants antigen residues.

Noncovalent contacts were derived from Arpeggio^53^ donor/acceptor interaction output generated for each complex. Eight interaction classes were analyzed: hydrogen bonds, hydrophobic contacts, van der Waals interactions, van der Waals clashes, weak polar contacts, polar contacts, aromatic interactions, and ionic interactions.

Interaction networks were constructed in NetworkX^54^ as bipartite graphs linking paratope to epitope residues. Nodes were annotated with residue number, residue name, residue role, *protein-quality*/*protein-interaction* classification, and paratope/epitope identity (primary versus control). Edges were weighted by total interaction count and colored according to the dominant interaction class. Node outlines encoded structural relationships. Networks were rendered using a left–right bipartite layout and assembled into multi-complex figure panels with unified visual styling and legend structure.

For comparative analysis, antigen residues were assigned to *protein-quality*, *protein-interaction*, or other groups. Interaction counts were normalized by the number of residues in each group to correct for differences in group size. We compared interaction density per residue, the fractional composition of interaction types, and absolute interaction rates normalized by residue counts, and we examined per-complex distributions of *protein-quality* and *protein-interaction* interactions. All statistical analyses were performed using SciPy, and visualizations were produced with seaborn using a consistent color scheme.

All biochemical analyses were performed in Python 3.10 using pandas^55^, joblib, NetworkX^54^, matplotlib^56^, seaborn^57^, and SciPy^58^.

## Data availability

All data underlying figures will be released upon approval of the manuscript.

## Code availability

The code for processing the data and creating the figures in this manuscript will be released upon approval.

## Author contributions

Conceived research: RS, VG

Supervised research: JGvB, SD, RS, VG

Designed wet-lab experiments: JdK, MA, YW, HB, EE, SD, RS

Performed wet-lab experiments: EE

Analytics: JdK, ES, AM, MA, LMB, MF, PR, HB, LiWo, YW, LuWe, LL, EE, RE

Interpretation: JdK, ES, AM, MA, LMB, LL, RE, RL, SD, RS, VG

Wrote manuscript: JdK, ES, RS, SD, VG

Edited manuscript: JdK, ES, AM, MA, LMB, MF, PR, HB, LiWo, YW, LuWe, LL, EE, KLM, RE, RL, JGvB, SD, RS, VG

## Acknowledgments

The authors would like to thank Fridtjof Lund-Johansen (University of Oslo) for critical review of this manuscript.

## Supplementary information

**Supplementary Figure 1.**
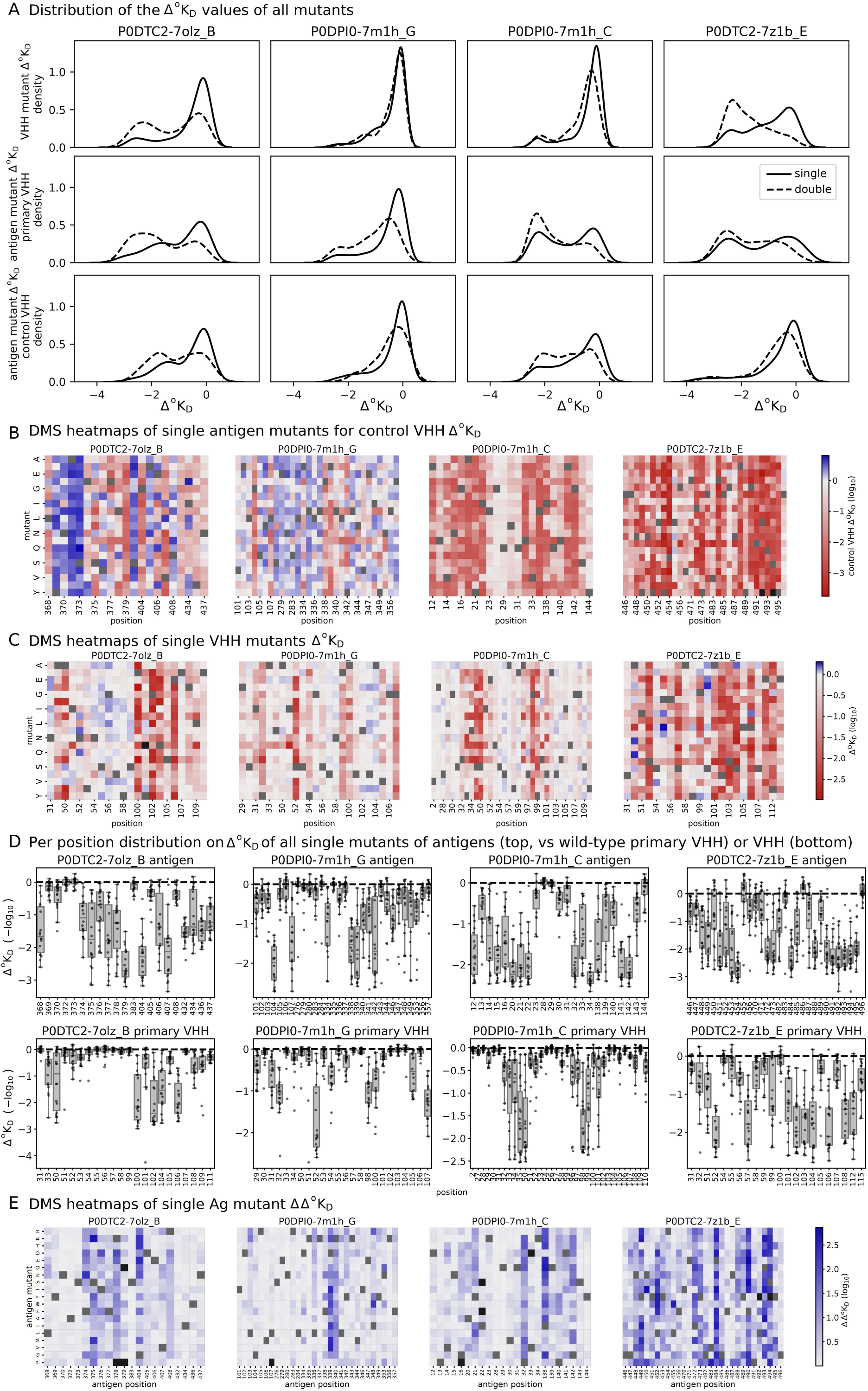
**The distributions of single epitope and paratope mutations in the four VHH-antigen complexes**. (A) The distribution of Δ°K_D_ values of the primary VHH (top) or control VHH (middle) of the single epitope mutations and the single primary VHH mutations (bottom) (B) Antigen DMS heatmaps, control VHH Δ°K_D_ (black is NaN, grey = WT). (C) Primary VHH DMS heatmaps (black is NaN, grey = WT). (D) The mutational tolerance per epitope (top) and paratope (bottom) position for the four complexes, as indicated by boxplots of Δ°K_D_. A large variability in mutational impact can be observed over different positions.

**Supplementary Figure 2.**
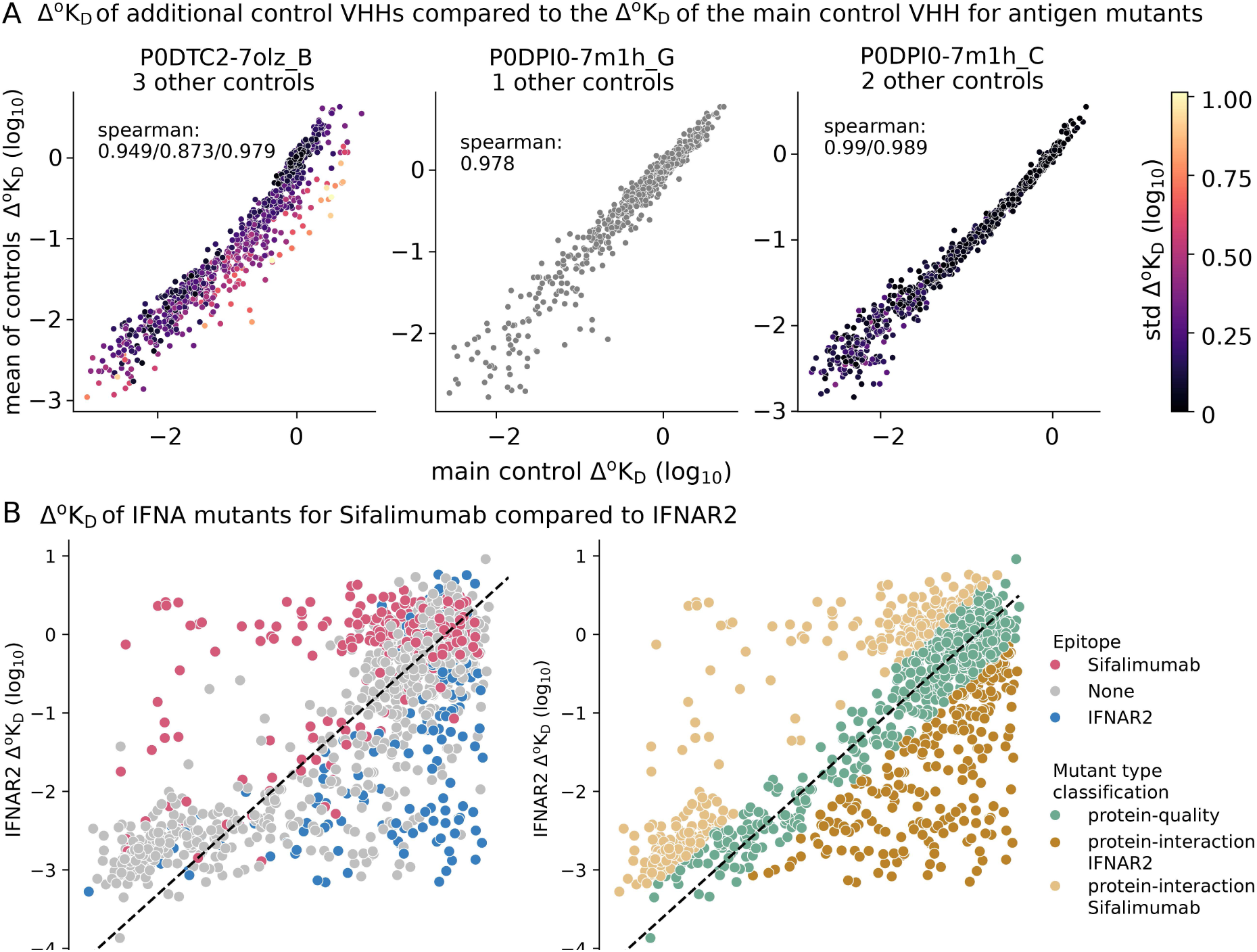
Protein quality and interaction changes in mutants bound by receptors and antibodies. (A) The correlation of the Δ°K_D_ of the main control VHHs on the x-axis, compared to the mean of the Δ°K_D_ of the other available controls on the y-axis. The spearman correlation of each of the separate controls with the main control is displayed. Each point is single mutant, and points are colored according to the std of the other controls (if more than 1). (B) The affinity of receptor IFNAR2 and Fab Sifalimumab for single mutants of IFNA. (left) the mutations are colored by the epitope of the corresponding binder (i.e., positions with any atom ≤5 angstrom distance to the binder). (right) the mutations are colored by the type of mutation. Protein-interaction variants are determined as ΔΔ°K_D_ > log_10_(5). (C) Similar to A, but for Fabs and VHHs binding SARS-CoV-2 spike RBD from a public AlphaSeq dataset. On the x-axis, one of the controls is displayed (randomly selected), and on the y-axis the mean of the other controls. Mutants are colored by the std of the controls.

**Supplementary Figure 3.**
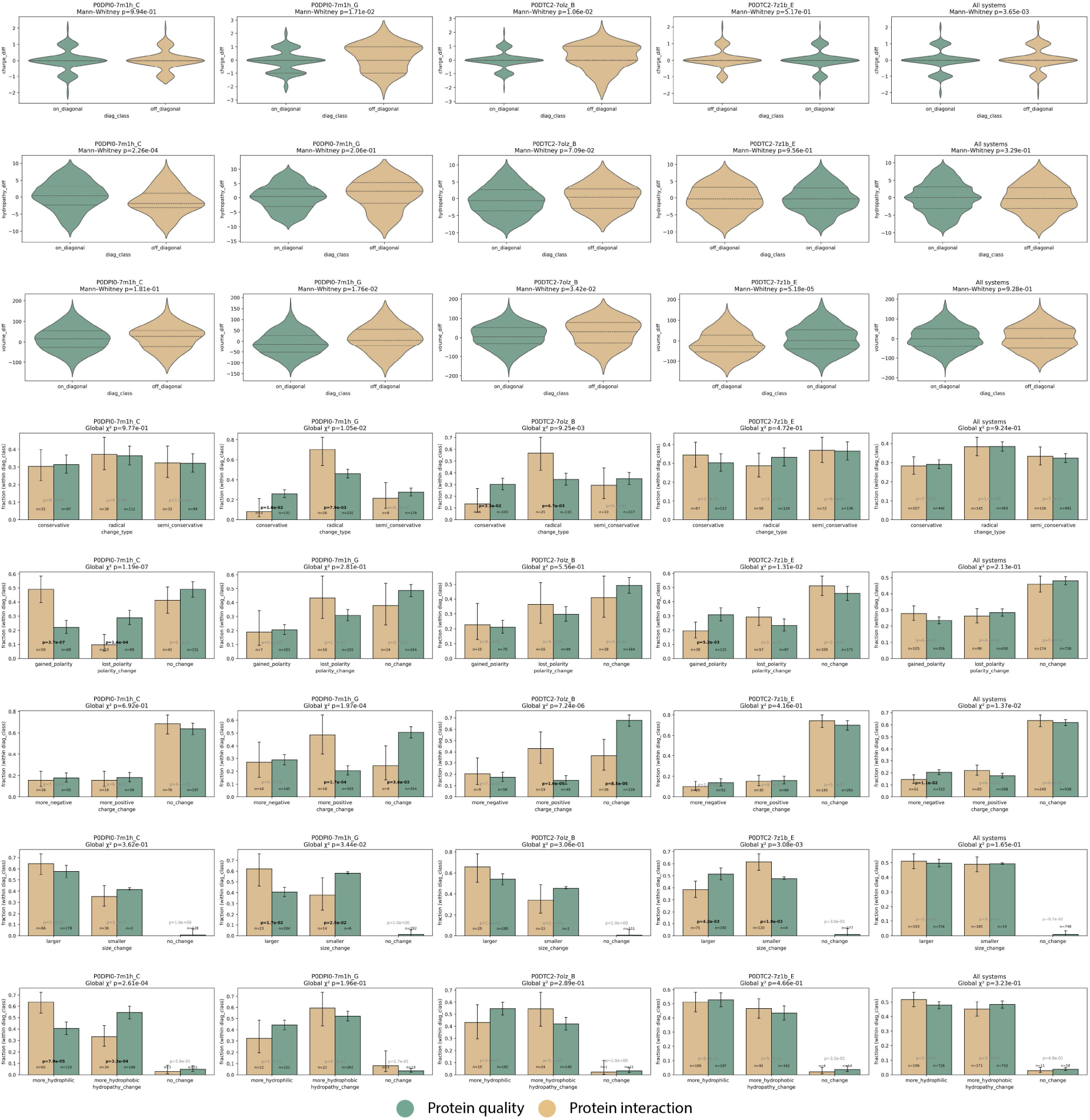
Biochemical changes upon single amino acid mutations at protein-quality and protein-interaction positions. Violin and bar plots summarizing the biochemical consequences of all single-amino-acid substitutions occurring at positions classified as protein-quality (green) or protein-interaction (gold). For each complex, mutations were annotated for amino-acid similarity class, polarity, charge, volume, and hydropathy changes. Violin plots show the distribution of continuous values, while bar plots show the frequency of categorical values. Error bars represent binomial 95% confidence intervals for categorical features and interquartile ranges for continuous features. These comparisons highlight systematic biochemical preferences associated with protein-interaction (protein-interaction) versus protein-quality (protein-quality) effects.

**Supplementary Figure 4.**
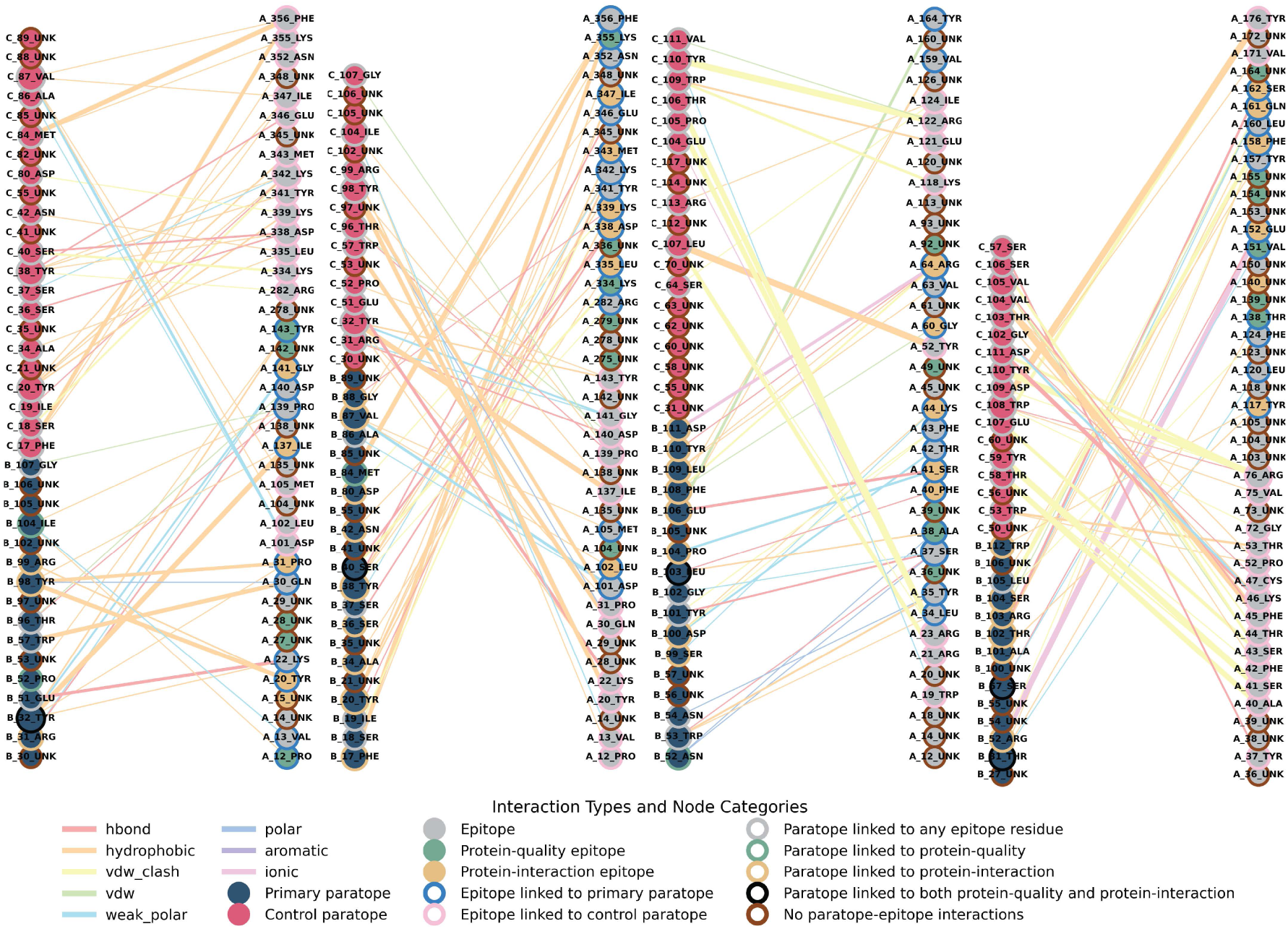
Residue-level antigen-paratope interaction networks as structural determinants of protein-quality and protein-interaction behavior. Bipartite residue-interaction networks for each antigen-VHH complex, linking epitope residues to primary and control paratope residues. Epitope and paratope definitions were assigned using a 4.5 Å distance threshold applied to post-processed structures curated for this study. Nodes represent individual residues: primary paratope residues are displayed in dark blue and control paratope are shown in pink, epitope residues are colored according to their functional classification, with protein-quality residues in green, protein-interaction residues in gold, and unclassified residues in grey. Node outlines encode their interaction context. For paratopes, the outline color reflects the type of epitope contacted: green if the residue interacts exclusively with protein-quality epitopes, gold if it interacts exclusively with protein-interaction epitopes, and black if it contacts both classes. For epitopes, the outline indicates the type of the contacting paratope: blue when contacting only primary VHH paratopes and pink when contacting only control VHH paratopes. Residues that form no paratope-epitope interactions in the wild-type complex (common among protein-quality epitope positions) are marked with a brown outline. Edges correspond to individual residue-residue interatomic contacts, colored according to the interaction type for that residue pair: red for hydrogen bonds, orange for hydrophobic contacts, yellow for van der Waals clashes, light green for van der Waals contacts, cyan for weak polar interactions, blue for polar interactions, purple for aromatic contacts, and pink for ionic interactions. Edge thickness scales with the total number of the specific type of interatomic interactions between the residue pair.

**Supplementary Figure 5.**
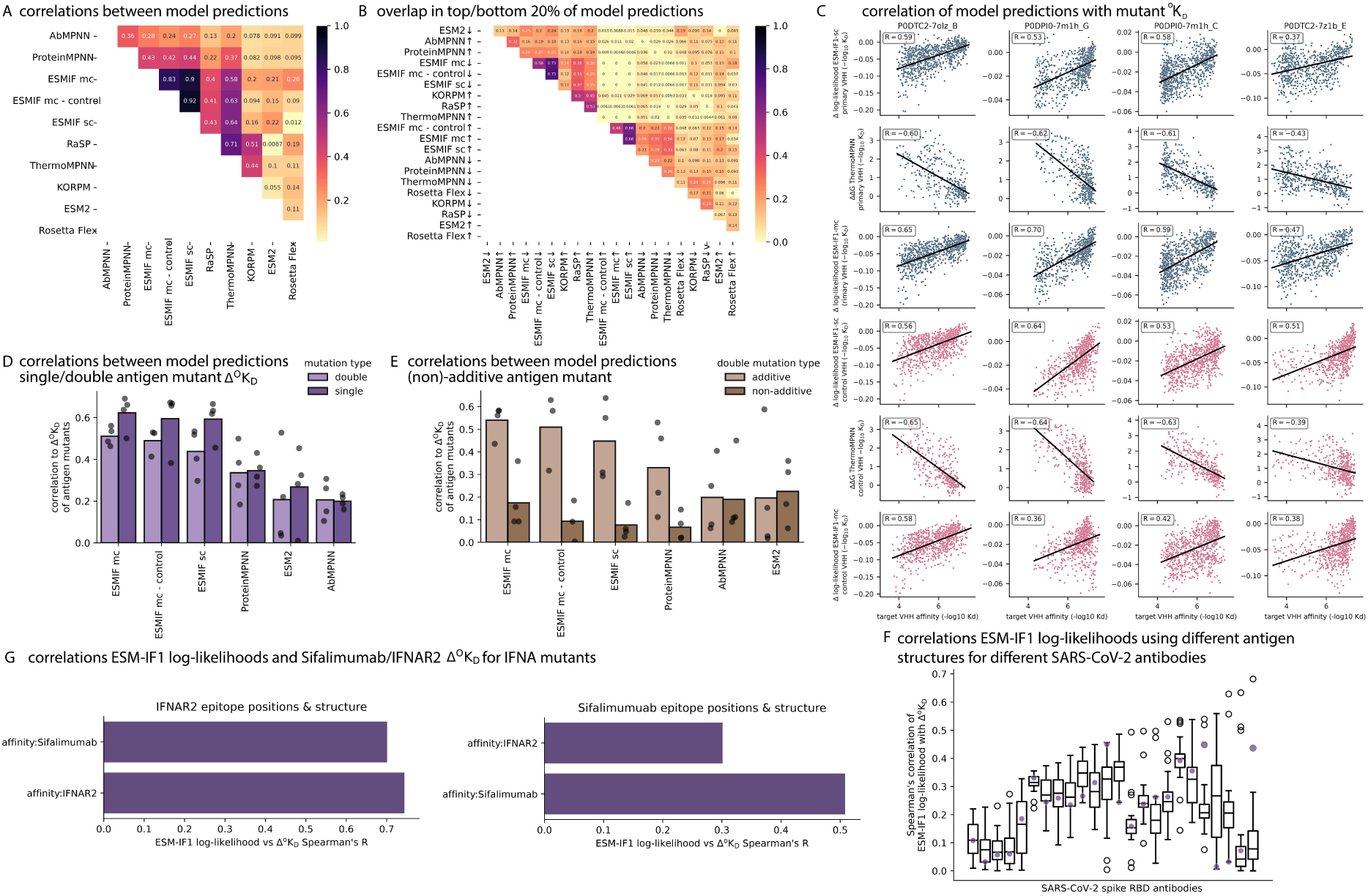
Existing models mostly predict protein-quality and not protein interaction. (A) The Spearman correlation between a variety of models’ predictive scores of single epitope mutations. The same models are shown as in Fig. 4A. (B) The overlap in the top (↑) or bottom (↓) 10% of predictions of different models. (C) The correlations of ESM-IF1 single-chain, ThermoMPNN, and ESM-IF1 multi-chain prediction scores for single epitope mutations with Δ°K_D_ for the four complexes. Each point is a mutant sequence. (D) The Spearman correlation between the Δ°K_D_ of the primary VHH of epitope single and double mutations and the prediction score of the models that can ingest double mutations. (E) Similar to D, but additive and non-additive. (F) ESM-IF1 single-chain log-likelihood correlation of IFNA single mutants to the Δ°K_D_ of Sifalimumab and IFNAR2. ESM-IF1 was conditioned on the antigen chain of the IFNAR2 structure and applied on IFNAR2 epitope positions (left), or on the antigen chain of the Sifalimumab structure and the Sifalimumab structure (right). (G) ESM-IF1 log-likelihoods were predicted 24 times for single SARS-CoV-2 mutants, each time using the antigen chain of a different RBD-antibody structure. The boxplots display the distribution of the correlation of these 24 sets of ESM-IF1 to the Δ°K_D_ of each antibody, based on the mutants of that antibody’s epitope. The purple dot indicates the correlation for when the antigen chain of the antibody’s own structure was used.

**Supplementary Figure 6.**
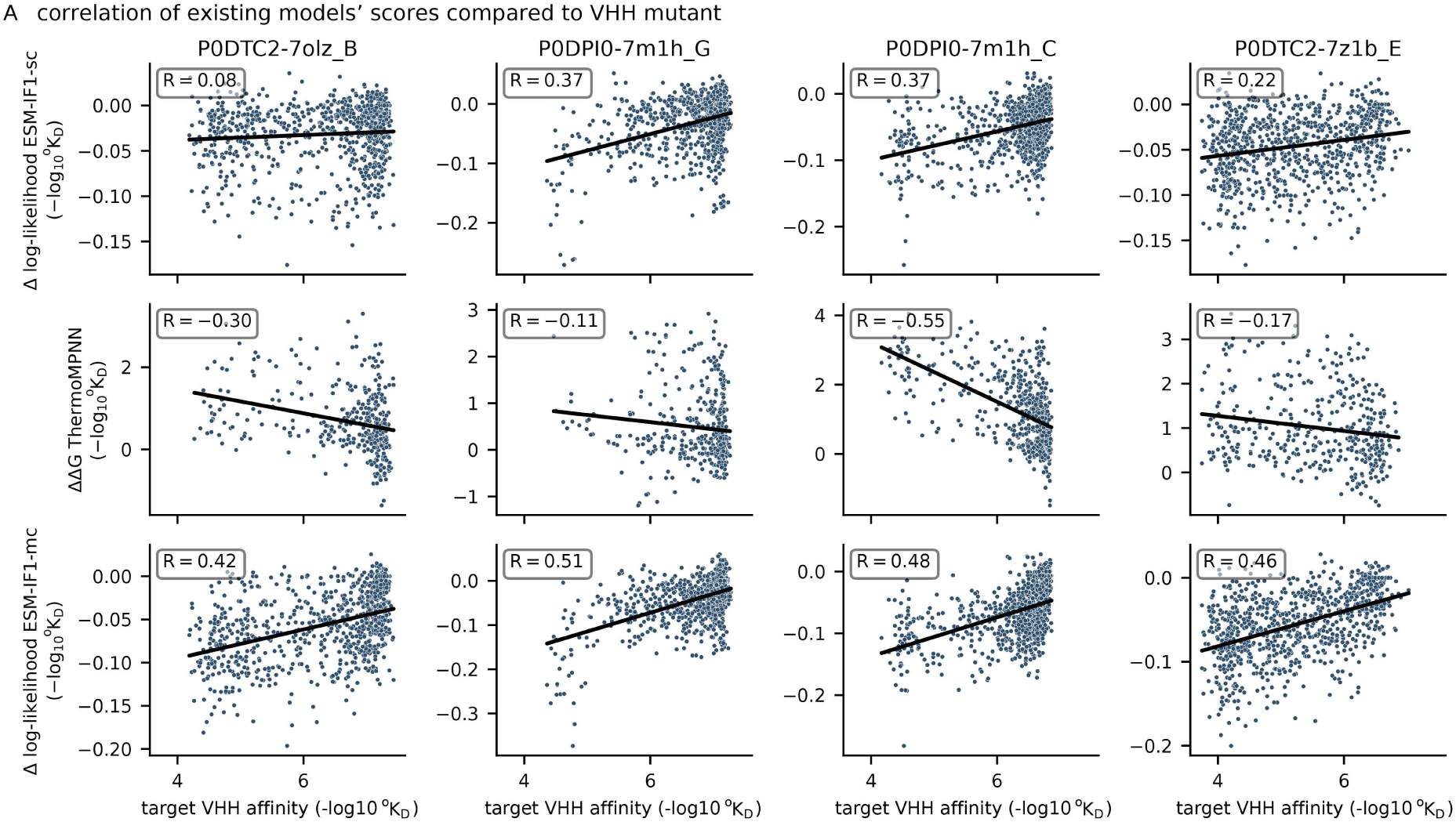
Models’ prediction scores correlate less with VHH mutations than Ag mutations. (A) The Spearman correlations of ESM-IF1 single-chain, ThermoMPNN, and ESM-IF1 multi-chain prediction scores for single paratope mutations with Δ°K_D_ for the four complexes. Each data point is a mutant sequence.

**Supplementary Figure 7.**
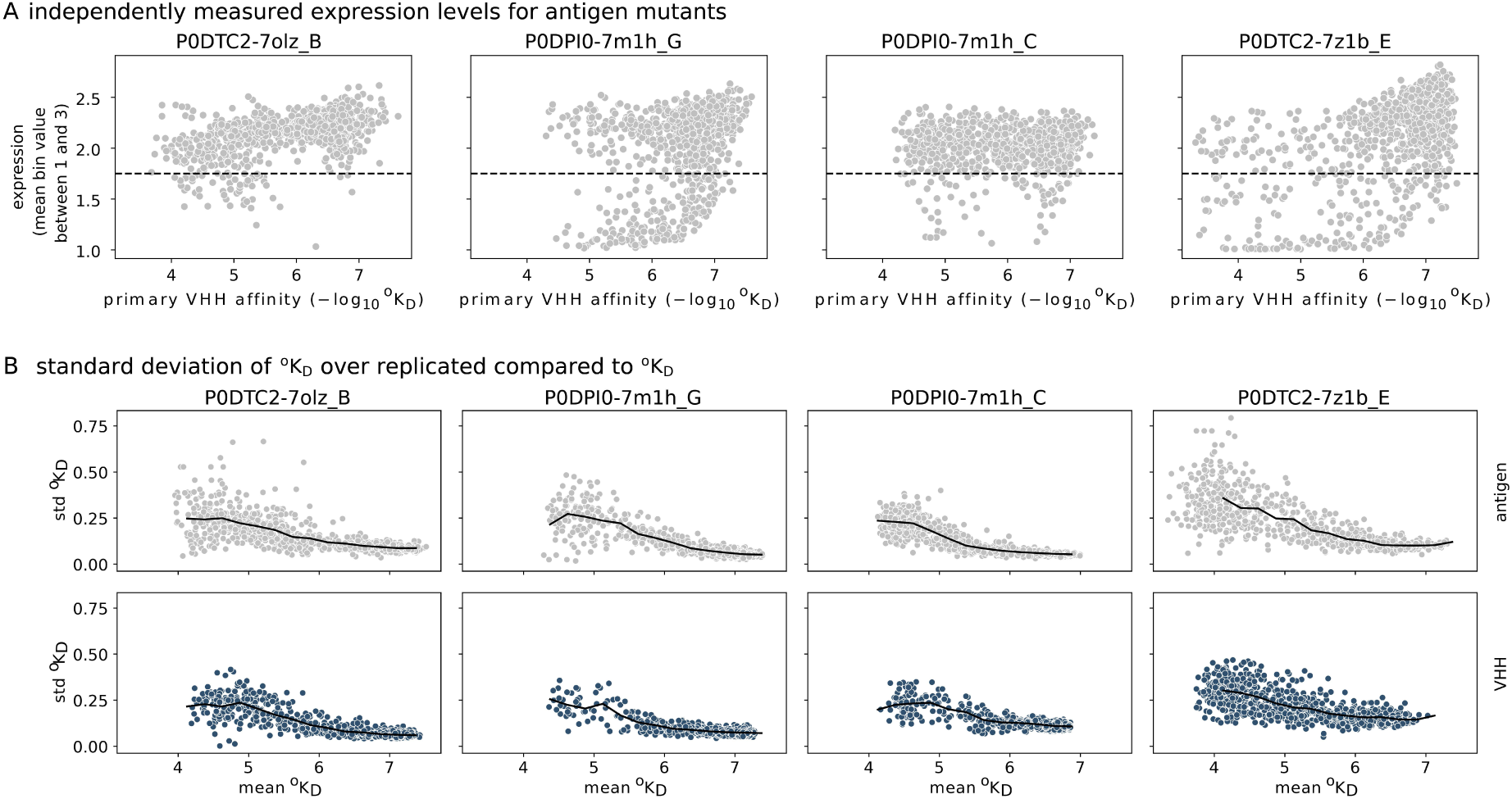
**Data patterns in AlphaSeq observed affinity. (**A) Expression measurement distribution compared to primary VHH observed affinity for all antigens. (B) the standard deviation of each antigen (top) or VHH (bottom) mutation of the four complexes over the 10 synonymous wild-type replicates of the other protein compared to the mean of those 10 measurements. The average standard deviation of values binned by the mean in bins of 0.25 °K_D_ on the x-axis are depicted by a black line. For low affinity measurements, the standard deviation is lower.

